# From nucleotides to semantics: genomic representation learning via joint-embedding predictive architecture

**DOI:** 10.64898/2026.04.02.716255

**Authors:** Chengsen Wang, Qi Qi, Haifeng Sun, Zirui Zhuang, Bo He, Siying Liu, Jianxin Liao, Jingyu Wang

**Author notes:** Contributing authors. These authors contributed equally to this work (Chengsen Wang, Qi Qi).

## Abstract

Decoding the regulatory syntax encoded in genomic sequences is a central objective in computational biology. Most existing genomic foundation models treat DNA as a language and adopt pretraining objectives from natural language processing. DNA sequences, however, lack explicit semantic boundaries and contain substantial evolutionary noise. Nucleotide-level reconstruction in a low-dimensional input space can therefore increase computational overhead and may yield representations with limited discriminative capacity. Downstream tasks often depend on expensive finetuning, which restricts practical use in many biology laboratories. Here we present GenoJEPA, a genomic representation learning framework based on joint-embedding predictive architecture. GenoJEPA combines continuous patching with semantic alignment, shifting the optimization from local base reconstruction to semantic alignment in latent space. Across 55 downstream tasks, GenoJEPA shows strong representational capacity and robust generalization while reducing parameter count and computational cost. The resulting semantic vectors from frozen GenoJEPA support lightweight GPU-free classifiers to achieve competitive accuracy. These results suggest a practical route towards efficient training and broad application of larger-scale genomic foundation models.

## 1 Introduction

Decoding the regulatory syntax embedded in DNA sequences and understanding evolutionary mechanisms are central goals of computational biology [1, 2]. Recently, unsupervised foundation models have advanced genomic research [3, 4]. By pretraining on large-scale sequencing data, these models capture cross-species genetic patterns and support prediction of biological processes such as gene expression and epigenetic modification. Most existing models, inspired by the success of Natural Language Processing (NLP), treat the genome as a form of biological language and use Masked Language Modelling (MLM) [5] or Next-Token Prediction (NTP) [6] during pretraining. Despite differences in tokenization [7, 8] and backbone design [9, 10], these approaches still rely on linguistic modelling. Important differences between genomic sequences and human language, however, limit this analogy.

Human language is an artificial system with high information density, strong syntactic priors, and explicit lexical boundaries [11]. Genomic sequences, although composed of discrete nucleotides, are organized more like natural images than like text. Both genomic sequences and images arise through natural processes, lack predefined semantic boundaries, and exhibit a relatively low signal-to-noise ratio [12, 13]. Applying frameworks developed for information-dense human language directly to genomic data may therefore hinder efficient extraction of semantic features from a background of evolutionary noise.

This mismatch affects both pretraining and downstream adaptation. Generative and reconstructive objectives require nucleotide-level recovery in a low-dimensional input space, much as Masked Autoencoders (MAE) [14] reconstruct pixel-level details in Computer Vision (CV). Such objectives can encourage models to spend capacity on fitting noise with limited regulatory relevance and may yield representations with limited discriminative power [12, 15]. Because high-frequency noise is not adequately filtered during pretraining, researchers often need extensive parameter updates for each downstream task, which require far more computation than inference with a frozen backbone. This dependence has become a practical bottleneck in computational biology, where model scale continues to grow but computing resources remain limited in many laboratories. Deeper architectures and longer context windows increase theoretical capacity [7, 16], yet they also raise the barrier to downstream use. For many biology laboratories and clinical research institutions, procuring and maintaining high-end computing infrastructure remains prohibitive. The field therefore needs pretrained encoders that can serve as reliable feature extractors while remaining frozen during downstream application.

Motivated by these challenges and the structural similarity between DNA sequences and natural images, we introduce an alternative framework for genomic sequence modelling. GenoJEPA is an unsupervised Joint-Embedding Predictive Architecture (JEPA) [12] framework that shifts optimization from nucleotide reconstruction to semantic alignment in a high-dimensional latent space. By aligning features in latent space, the model can suppress polymorphic variation and devote more capacity to biologically relevant structure, yielding transferable representations for downstream tasks without extensive finetuning [17]. Furthermore, at the input stage, we replace conventional genomic tokenization with a continuous patching strategy [18] widely used in CV. Common tokenization schemes are based on Byte-Pair Encoding (BPE) [19] or fixed-window k-mers [20]. Both can introduce vocabulary redundancy and are sensitive to local mutations. Continuous patching instead treats genomic segments as local signals and maps them directly into a dense vector space, which avoids discrete vocabulary inflation while preserving biochemical dependencies within the sequence. After pretraining on a multispecies genomic corpus spanning 850 representative species across major taxonomic lineages [7], GenoJEPA learns transferable representations that perform well across diverse downstream tasks.

We evaluated GenoJEPA against five representative baselines on 55 tasks from three established genomic benchmarks [7, 19, 21]. These tasks cover regulatory element identification, epigenetic mark prediction, splice site recognition, and cross-species sequence classification, among others. The evaluation includes probing, finetuning, few-shot, and efficiency analyses rather than a single downstream scenario.

In terms of parameter efficiency, GenoJEPA achieves competitive or stronger recognition performance with only one-tenth to one-hundredth of the parameters of conventional Transformer baselines [7, 19]. Relative to emerging architectures designed for long-sequence processing [10, 22], it also shows shorter running times and more stable memory usage in most evaluated settings. GenoJEPA further shows favourable data efficiency. With only a small fraction of the available training data, it approaches the performance of strongbaselines trained on the complete dataset. These findings suggest that GenoJEPA is well suited to resource-limited and annotation-scarce biological settings.

GenoJEPA also performs strongly under both finetuning and probing protocols. In a comparison with NT-v2, which was pretrained on the same 850-species corpus with nearly 10 times more parameters, GenoJEPA learned stronger frozen representations and remained slightly better under finetuning. Strong finetuning performance supports its suitability in accuracy-sensitive settings. The improvement in probing may be especially relevant in practice because it indicates that GenoJEPA can provide discriminative representations without finetuning. This property allows researchers with limited computational resources to achieve competitive downstream performance using lightweight machine learning models without GPU acceleration. Overall, these results suggest that latent-space semantic alignment offers a practical route towards more efficient training and broader application of larger-scale foundation models in genomic research.

## 2 Results

### 2.1 GenoJEPA learns genomic representations via joint-embedding prediction

Genomic sequences encode complex evolutionary patterns and regulatory logic. Representing these sequences as continuous features suitable for computation remains a central challenge in computational biology. GenoJEPA is a self-supervised framework that adapts the LeJEPA [23] formulation to genomic sequences, captures biological semantics in latent space, and provides transferable genomic representations.

As shown in Figure 1a, the model begins by tokenizing the input DNA sequence. Mainstream BPE [19] relies on frequency statistics that do not necessarily reflect functional conservation. K-mer segmentation [7] may split biologically meaningful motifs at arbitrary boundaries, and large values of *k* lead to impractically large vocabularies. These discrete schemes are also sensitive to sequence variation. A single-nucleotide mutation can map highly homologous segments to different identifiers and obscure their evolutionary relationship. Single-nucleotide tokenization [22] alleviates some of these issues, but its computational cost grows quadratically with sequence length. Stacked convolutional layers [24] can reduce this cost through local downsampling, yet they also consume a substantial share of the backbone parameter budget. GenoJEPA instead partitions each input sequence into non-overlapping nucleotide patches [18] and maps them to continuous feature vectors through linear projection. This design avoids vocabulary inflation, shortens the effective sequence length, and preserves local biochemical dependencies.

**Fig. 1.**
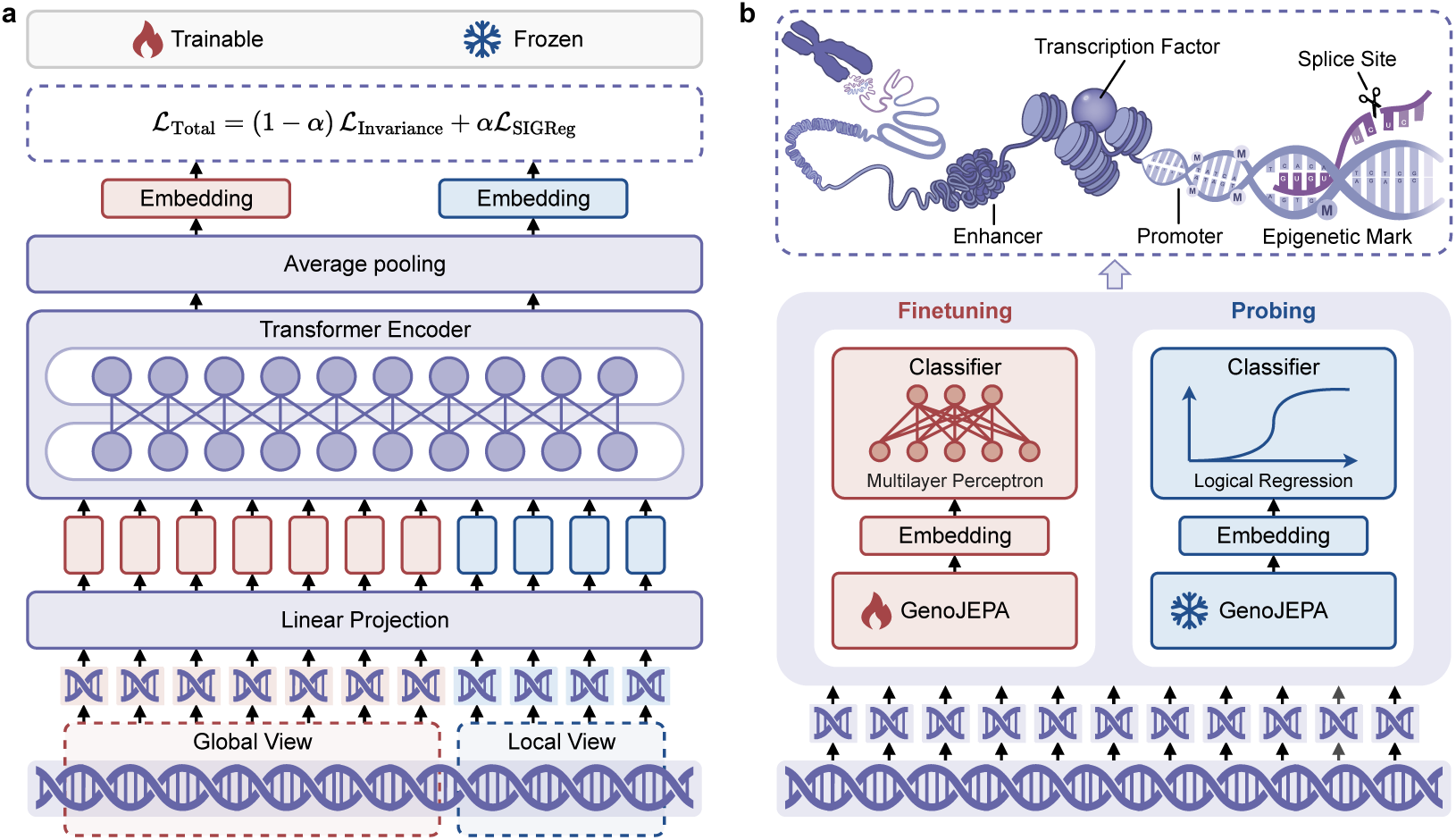
GenoJEPA learns genomic representations via joint-embedding prediction. **(a)** shows the pretraining framework. The input DNA sequence is split into non-overlapping patches, linearly projected into dense embeddings, and processed by a Transformer encoder. Token embeddings are then averaged to obtain a sequence representation. Following the LeJEPA formulation, the sequence is augmented into multiple local and global views through random cropping. GenoJEPA is optimized with an invariance loss that aligns semantic features across views together with a SIGReg loss that prevents representation collapse. **(b)** shows the downstream adaptation strategies for genomic tasks such as functional element identification. In resourcelimited settings, the pretrained encoder is frozen and the resulting sequence embeddings are passed to a lightweight task head for probing. In accuracy-sensitive settings, the entire GenoJEPA model is finetuned end to end together with a task head.

In the modelling stage, these dense vectors are passed to a Transformer encoder based on the ModernBERT [25] architecture. This backbone retains the advantages of bidirectional encoders for semantic extraction while incorporating design advances from recent autoregressive Large Language Models (LLM) [26–28]. Token representations are then averaged to obtain a sequence representation. In line with the multi-view framework used in the JEPA family [17, 23], each sequence is transformed into multiple local and global views through data augmentation such as random cropping. GenoJEPA follows the LeJEPA [23] objective and aligns all views to the mean representation of the global views in a high-dimensional latent space rather than reconstructing the original low-dimensional nucleotide sequence. To maintain feature discriminability and prevent representation collapse, training further includes a Sketched Isotropic Gaussian Regularization (SIGReg) [23] loss, which guides the learned features towards a distribution that supports discrimination. During pretraining, a nonlinear projection head maps sequence representations into a dedicated latent space for loss computation. This helps preserve the generality of the backbone and reduces overfitting to the pretraining objective [29].

To capture broad evolutionary constraints and underlying regulatory syntax, the pretraining corpus aggregated genomic data from 850 representative species [7] and comprised nearly 200 billion nucleotides after quality filtering. Owing to continuous patching and latent-space feature alignment, GenoJEPA reduced the overhead of long-sequence processing and local detail reconstruction, which supported favourable training efficiency. We instantiated the framework at two scales, a lightweight version named GenoJEPA-T with 6M parameters and a base version named GenoJEPA-B with 52M parameters. GenoJEPA-T required only a single RTX 3090 24 GB GPU and completed 100,000 training steps in about 12 hours. GenoJEPA-B completed 500,000 pretraining steps on a single A800 80 GB GPU in about 150 hours. This training setup balanced computational accessibility and data throughput and provided a practical basis for future scaling.

For downstream genomic tasks, Figure 1b shows two complementary modes of use. In laboratories with limited computational resources, researchers can freeze the pretrained backbone and use probing, which gives a direct view of native representation quality. In this setting, the input DNA sequence passes through GenoJEPA to produce token embeddings that are aggregated into a sequence representation. A lightweight classifier such as logistic regression [30] can then be trained on these static features. This setup is well suited to workflows in which feature extraction is performed on remote infrastructure and downstream analysis is handled locally. In settings with stringent accuracy requirements, the model can instead be finetuned end to end. The backbone and an attached nonlinear classifier implemented as a Multi-Layer Perceptron (MLP) are jointly updated, allowing the pretrained model to adapt more fully to the target distribution.

### 2.2 GenoJEPA maintains highly discriminative features under probing evaluation

Current evaluations [7, 19] of genomic foundation models often rely on task-specific finetuning. This practice can bias model comparison and make it difficult to attribute performance differences directly to pretrained representations. For example, updating different layers during finetuning can lead to different degrees of overfitting [31]. Parameter-efficient finetuning methods [32] add further variation through extra hyperparameters. To assess the representations learned during pretraining, we froze all backbone parameters and trained only a lightweight classifier for probing. Compared with finetuning, this frozen-backbone setup offers a more direct comparison of representation quality across models.

The experiments covered three widely used evaluation benchmarks, Genomic Benchmarks (Genomic) [21], GUE Benchmarks (GUE) [19], and Nucleotide Transformer Tasks (NT Tasks) [7]. Together they contain 55 subtasks spanning cross-species sequence classification, promoter and enhancer detection, RNA splice site recognition, and transcription factor binding prediction. For comparison, we included several representative genomic foundation models, HyenaDNA (7M) [22], CaduceusPh (8M) [10], GROVER (87M) [8], DNABERT-2 (117M) [19], and NT-v2 (494M) [7]. These models cover different tokenization strategies and backbone architectures. NT-v2 is particularly informative because it was pretrained on the same 850-species corpus as GenoJEPA while using nearly 10 times more parameters. All baselines used their optimal official weights pretrained with MLM or NTP.

For sequence feature extraction, we applied average pooling to the token representations. Prior work [31] has shown that this aggregation step provides stable behaviour. We used a logistic regression [30] classifier with the default scikit-learn configuration [33] because it is efficient, GPU-free, and introduces little hyperparameter overhead. Because many genomic datasets are imbalanced, we adopted the Matthews Correlation Coefficient (MCC) [34] as the primary metric, following established practice [7, 19]. All tasks were evaluated with 10fold cross-validation with a held-out test set. Detailed probing results for all models are provided in Table S28 in the Appendix.

Under this unified probing protocol, GenoJEPA-B achieved the strongest overall performance. Figure 2a shows that GenoJEPA-B won more tasks than any baseline under the pairwise Wilcoxon signed-rank test at *p* = 0.05. The lightweight GenoJEPA-T also remained competitive with NT-v2, which has roughly 100 times more parameters. Because GenoJEPA and NT-v2 were pretrained on the same 850-species corpus, broader data coverage is a less likely explanation for this result. The comparison is instead consistent with the view that the pretraining objective and the resulting representation geometry are important determinants of downstream transfer.

**Fig. 2.**
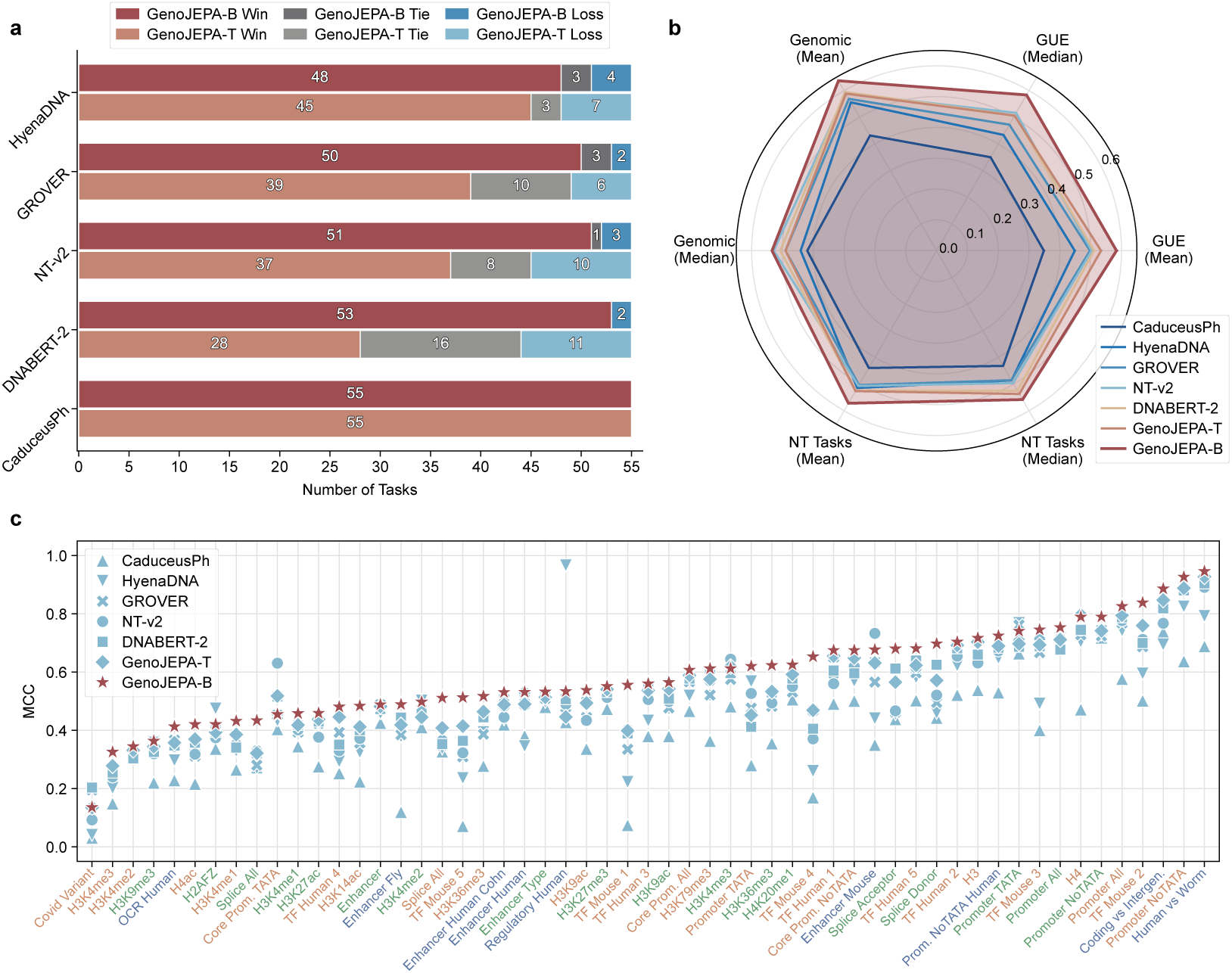
GenoJEPA maintains highly discriminative features under probing evaluation. Each task is evaluated through 10-fold cross-validation, and the mean test score is used as the representative result. **(a)** shows the probing win-loss comparison between GenoJEPA and the baselines across all benchmarks. A paired Wilcoxon signed-rank test at *p* = 0.05 is applied to the fold-wise test scores as a stability-aware paired comparison. The number of wins, ties, and losses is reported across 55 tasks from three benchmarks. **(b)** shows probing performance for each benchmark. The average and median performance across all tasks within each benchmark are reported from the representative scores. **(c)** shows task-level probing performance. On the horizontal axis, blue labels denote the Genomic Benchmarks, red labels denote the GUE Benchmarks, and green labels denote the Nucleotide Transformer Tasks.

Figure 2b and Figure 2c show how these gains are distributed across benchmarks and tasks. In the benchmark-level summaries, GenoJEPA-B consistently matched or exceeded the overall performance profiles of the other models. The task-level plots show particularly strong performance on mouse transcription factor binding prediction, splice-donor recognition, promoter detection, and several histone-modification tasks. These processes often depend on coordinated cis-regulatory elements arranged in specific spatial patterns. The observed pattern is consistent with the view that continuous nucleotide segments combined with latent-space alignment across views help capture structural motifs that lack explicit boundaries but still carry functional associations. One exception is human regulatory element classification, where HyenaDNA performed well, which suggests that specialized architectures can retain advantages on some tasks.

Taken together, these results support modelling DNA sequences through semantic alignment in a high-dimensional latent space. The representation space learned by GenoJEPA yields discriminative features without requiring complex downstream transformations. Because this probing setup also requires minimal computation, it offers a practical option for laboratories with limited resources and supports workflows that combine centralized feature extraction with lightweight local analysis.

### 2.3 GenoJEPA achieves competitive performance with end-to-end finetuning

Although probing gives a direct assessment of representation quality, some applications require the highest possible task-specific accuracy, for example in focused functional genomics studies or rare pathogenic variant identification. Finetuning is therefore required to estimate performance after full downstream adaptation.

The data splits and evaluation metrics for finetuning were the same as those used for probing in Section 2.2, which enabled direct comparison across the two settings. For each baseline, we followed the pooling and classifier choices recommended in the original publication so that each model could operate close to its strongest practical adaptation regime. For GenoJEPA, we used the first classification token as the global sequence representation and attached a two-layer MLP with GELU [35] activation. To give each model a fair opportunity to adapt, we tuned batch size and learning rate within a predefined grid and reported the best converged performance. As in probing, all finetuning tasks were evaluated with 10fold cross-validation with a held-out test set. Detailed results for all models are presented in Table S29 in the Appendix.

Across 55 subtasks, GenoJEPA remained strong under finetuning. Figure 3a compares the average MCC of each model under probing and finetuning. GenoJEPA-B, with 52M parameters, improved average finetuning performance by 2.9% relative to NT-v2, which was pretrained on the same 850-species corpus with nearly 10 times more parameters, while the gain in probing was about 13.5%. GenoJEPA-T, with only 6M parameters, also remained competitive across model scales. In probing, it outperformed the other baselines, including NT-v2, which contains nearly 100 times more parameters. In finetuning, it remained stronger than similarly sized baselines such as HyenaDNA and CaduceusPh and also outperformed GROVER, which has more than 10 times more parameters. The average task ranks in Figures 3b and 3c provide a view that is less sensitive to task difficulty. In probing, the average rank favoured GenoJEPA-B. In finetuning, performance gaps narrowed after weight adaptation, yet GenoJEPA-B still retained the highest average rank despite its modest parameter count.

**Fig. 3.**
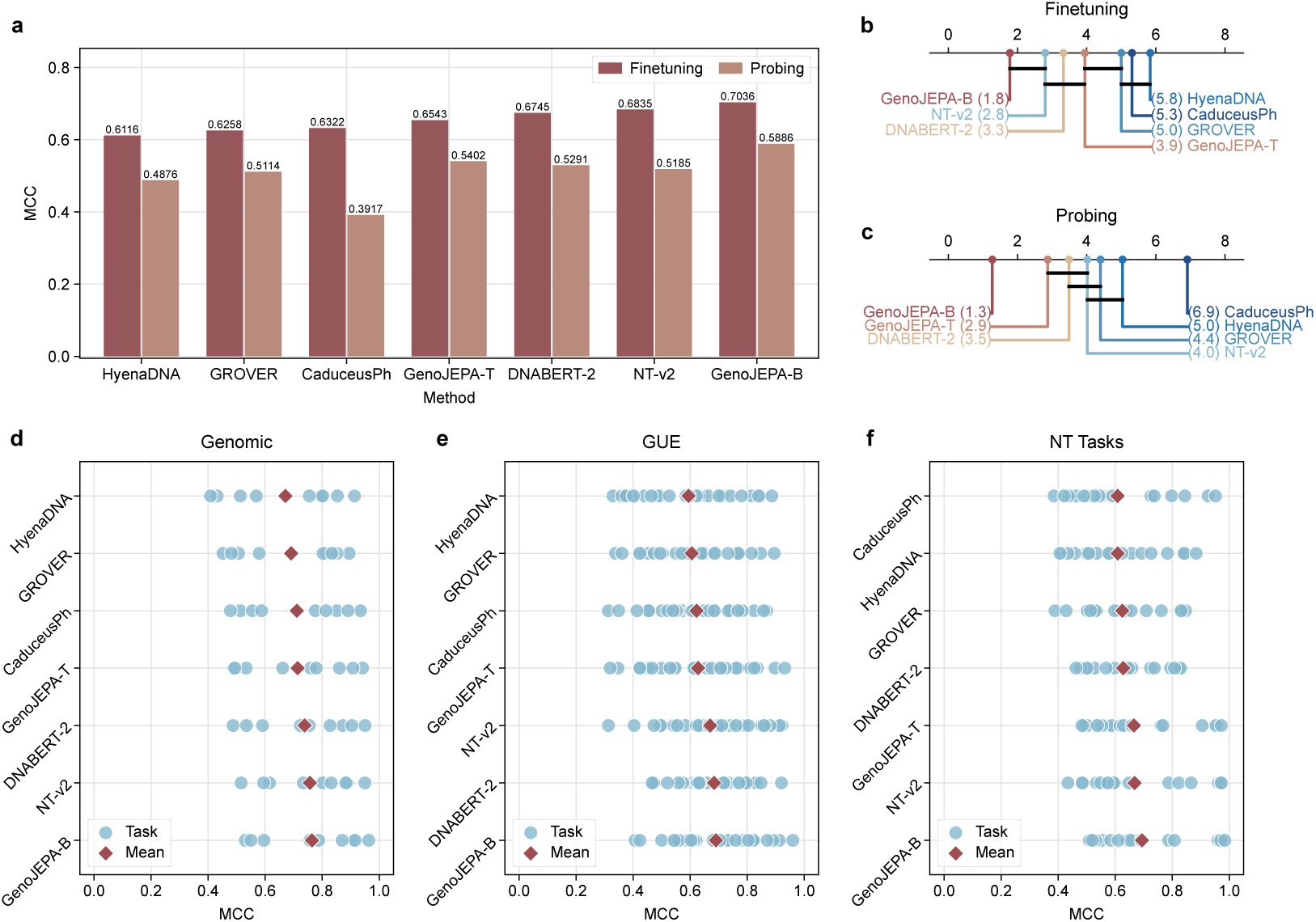
GenoJEPA achieves competitive performance with end-to-end finetuning. Each task is evaluated through 10-fold cross-validation, and the mean test score is used as the representative result. **(a)** shows the overall performance of GenoJEPA and the baselines across all benchmarks. The average performance across 55 tasks from three benchmarks is reported from these mean scores. **(b)** and **(c)** show the critical difference diagrams for finetuning and probing performance across all benchmarks. Statistical significance is summarized with the Friedman test at *p* = 0.05 followed by the Nemenyi post-hoc test at *p* = 0.05 on the task-level representative scores. The average rank and significance are reported from these mean scores. Black horizontal lines indicate no significant difference between methods. **(d)**, **(e)**, and **(f)** show finetuning performance for each task within the three benchmarks. Blue circles denote the mean score for each task, and red diamonds denote the average performance across all tasks within each benchmark.

A closer look at individual benchmarks provides further insight into the kinds of biological signals captured by GenoJEPA. As shown in Figure 3d, GenoJEPA-B remained among the stronger models on the Genomic Benchmarks, which include multispecies classification and regulatory element identification. Figure 3e shows the GUE results, where GenoJEPA scores clustered in the high-performance range across transcription-factor-binding and promoter-related tasks. On the Nucleotide Transformer Tasks in Figure 3f, GenoJEPA-B also remained competitive across human epigenetic and splice-related tasks. Some tasks remained difficult, including H4ac. Even so, the broad agreement between the probing and finetuning summaries points to consistent overall strengths.

Taken together with the probing results, this finetuning comparison is consistent with the view that replacing point-wise nucleotide reconstruction in a low-dimensional discrete space with alignment of continuous semantic features in a high-dimensional latent space can simplify downstream optimization while preserving core biophysical dependencies. Under the adaptation protocol used here, these initial weights allowed comparatively small models to approach or exceed much larger baselines.

### 2.4 GenoJEPA demonstrates favourable computational and memory efficiency

To provide practical guidance on balancing computational performance and resource use across downstream applications, we conducted a systematic efficiency analysis of each foundation model under controlled conditions, following the protocol in [31]. Runtime and memory are important bottlenecks in both research and clinical use, so we assessed both metrics.

To ensure a fair comparison, all experiments were conducted on a single A800 80 GB GPU with a batch size of 1 under the conditions used in [31]. Sequence lengths ranged from short fragments to beyond the hardware limit. To reduce stochastic noise and remove cold-start effects, each experiment was repeated for 10 epochs. The first 10 warmup steps of each epoch were discarded, and the average cost over the next 100 steps was reported as the final metric. To avoid bias from different sequence compression ratios introduced by different tokenization schemes, we measured length in native nucleotides. This common scale improves fairness across architectures and matches how researchers handle DNA fragments in practice. The trends are shown in Figure 4, with the left panel comparing models below 10M parameters and the right panel comparing models above 50M parameters.

**Fig. 4.**
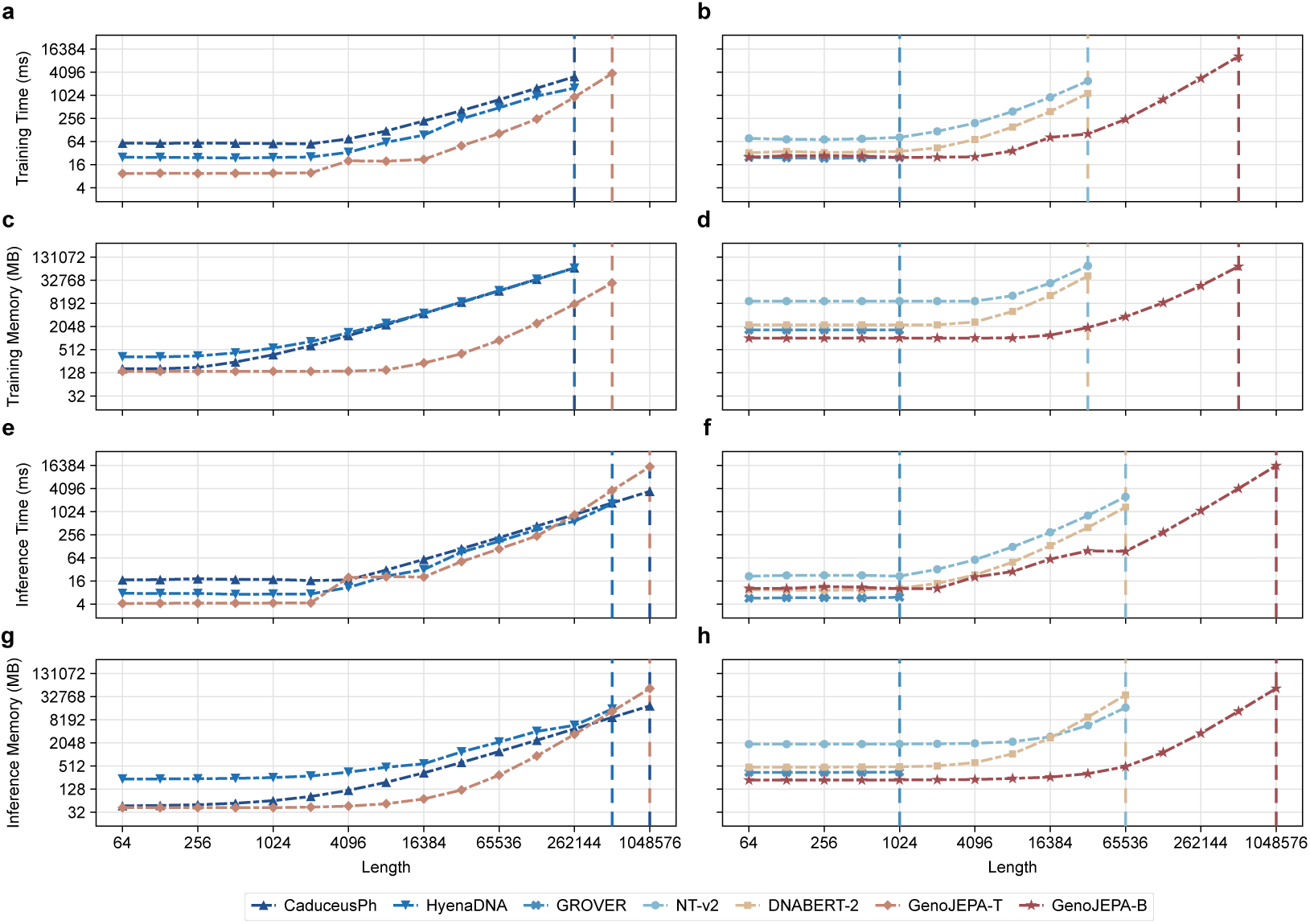
GenoJEPA demonstrates favourable computational and memory efficiency. **(a)**, **(c)**, **(e)**, and **(g)** report training time in ms, training memory in MB, inference time in ms, and inference memory in MB, respectively, for GenoJEPA and baselines with fewer than 10M parameters across different sequence lengths. **(b)**, **(d)**, **(f)**, and **(h)** report the same metrics for GenoJEPA and baselines with more than 50M parameters. Experiments are conducted with a batch size of 1 on an A800 80 GB GPU. Each experiment is repeated for 10 epochs, and mean values are reported. In each epoch, the first 10 steps are excluded to remove cold-start effects. Average metrics are then computed over the next 100 steps. Both axes are shown on a logarithmic scale. Vertical dashed lines indicate sequence lengths that exceed model support or require memory beyond GPU capacity.

For the larger models in the right panel, GenoJEPA-B shows lower runtime and memory use during both training and inference than DNABERT-2, GROVER, and NT-v2. This behaviour largely reflects the sequence compression produced by patching together with the relatively compact parameter count. GROVER achieves slightly lower inference time on shorter sequences, but its effective limit is capped at 510 tokens because it uses absolute position embeddings.

A more detailed view reveals a notable pattern in current sequence modelling. CaduceusPh and HyenaDNA are built on the Mamba [36] and Hyena [37] architectures, respectively, both of which are designed for sub-quadratic long-sequence processing. In practice, however, their runtime and memory use departed from these theoretical expectations. As shown in the left panel, when compared with GenoJEPA-T at the same sub-10M parameter scale, neither baseline showed the expected advantage in overall training or inference cost. The training and inference memory use of GenoJEPA-T remained stable until sequence lengths reached 8,192. CaduceusPh and HyenaDNA began to show increasing memory demands at training lengths of 256 and inference lengths of 1,024. During training, this earlier memory growth caused both models to encounter out-of-memory errors before GenoJEPA-T. During inference, GenoJEPA-T maintained a consistent overhead advantage across short and medium sequence lengths. Only when sequence lengths exceeded 262,144 did GenoJEPA-T become slower than CaduceusPh, which is expected from the quadratic cost of self-attention [38]. HyenaDNA did not show the expected long-sequence advantage and still encountered the out-of-memory errors at relatively short lengths.

This gap between theoretical complexity and practical efficiency becomes more pronounced in cross-scale comparisons. Here the comparison also includes GenoJEPA-B, which has nearly 10 times the parameters of the two baselines discussed above. Although its larger parameter count yields a somewhat higher initial memory footprint at very short sequence lengths, its training and inference memory use remained stable up to lengths of 16,384. During training, CaduceusPh and HyenaDNA did not show the expected speed advantage, and their runtime savings were limited. They also encountered out-of-memory errors earlier than GenoJEPA-B despite having only one-tenth as many parameters. Similar trends were observed during inference. In practice, these two baselines therefore face a tension between resource-intensive execution and limited probing performance.

In summary, GenoJEPA shows favourable efficiency relative to conventional large Transformer baselines and also compares well with newer architectures designed for efficient computation. Although CaduceusPh and HyenaDNA use architectures that are nominally more computation friendly, both incurred higher training and inference costs than GenoJEPA-T at comparable parameter counts across most tested ranges. Their training cost even exceeded that of GenoJEPA-B, which has about 10 times more parameters. Only the Mamba-based CaduceusPh retained a modest inference advantage on ultra-long sequences. The stability of GenoJEPA across the tested range suggests useful headroom for extending the model to longer genomic contexts in future.

### 2.5 GenoJEPA exhibits strong few-shot capabilities and feature versatility

Data efficiency is another practical criterion for foundation models. We therefore conducted few-shot probing evaluations within a framework that largely matched Section 2.2. The main difference was the training-data subsampling strategy. Within each fold, we performed stratified sampling at proportions from 10% to 50% for each class in the fitting and validation subsets while keeping the held-out test set unchanged. The same downsampled splits were used for all methods. Because the evaluation distribution was fixed, this comparison isolates reduced label availability from changes in test composition. Performance was then aggregated across 55 tasks from three benchmarks. As shown in Figure 5a, GenoJEPA maintained high discriminability across all data fractions. Its performance curve lay above those of the other baselines in the aggregated comparison. With only 10% of the training data, GenoJEPA reached a score of 0.5192, which was close to the best baseline score of 0.5253 at 50% and 0.5291 at full scale. This behaviour suggests that aligning continuous semantic features in high-dimensional space can reduce the dependence of downstream tasks on large labelled datasets.

**Fig. 5.**
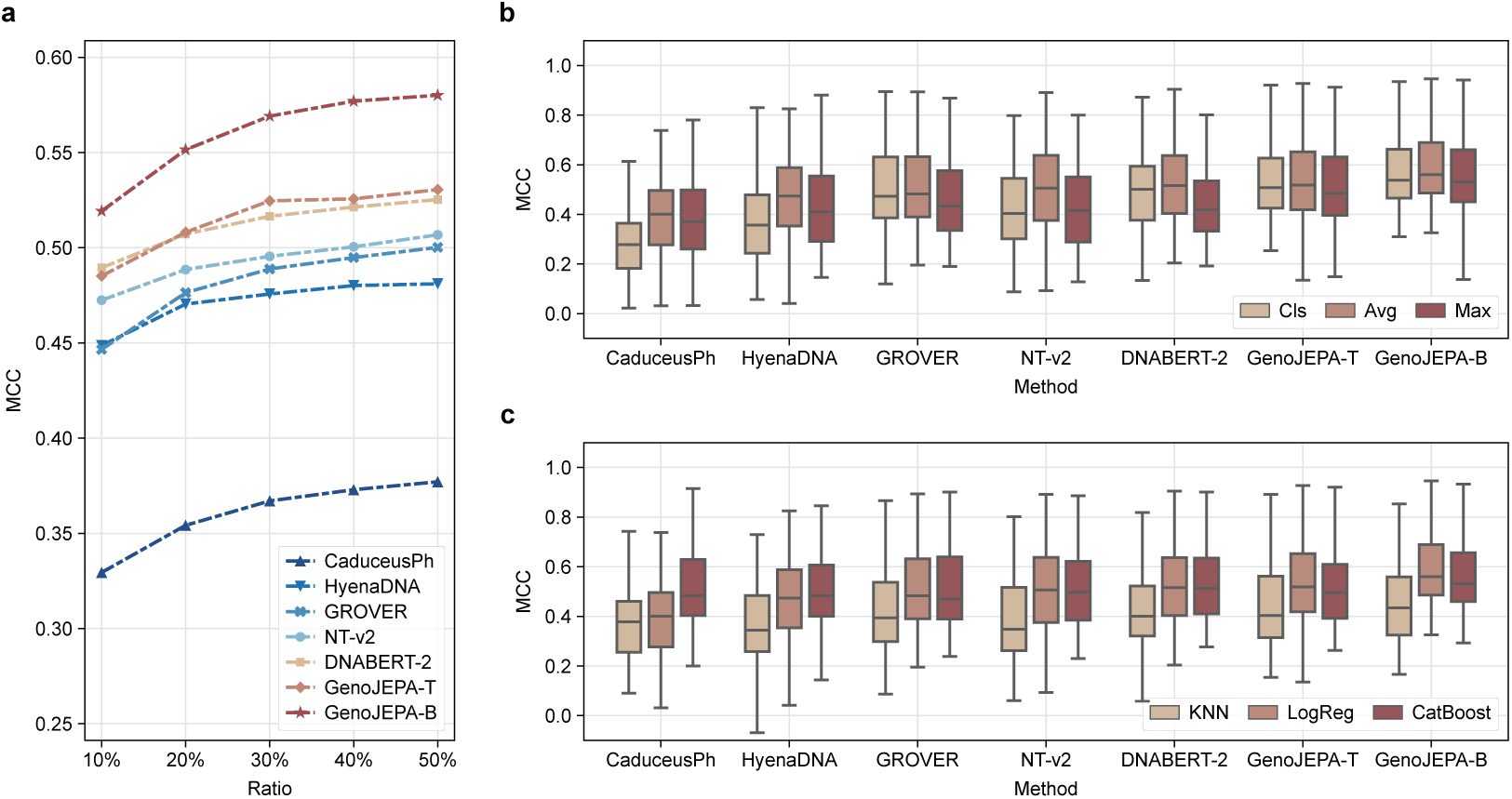
GenoJEPA exhibits strong few-shot capabilities and feature versatility. Each task is evaluated through 10-fold cross-validation, and the mean test score is used as the representative result. **(a)** shows probing performance for GenoJEPA and the baselines at different training-data scales across all benchmarks. Within each fold, the training and validation sets are stratified at proportions from 10% to 50% for each label while the test set is kept unchanged. Logistic regression with average pooling is used to assess the data efficiency of each method. The average performance across 55 tasks from three benchmarks is reported from these mean scores. **(b)** and **(c)** show probing performance under different pooling strategies and classifiers across all benchmarks. The boxplots summarize the distribution of representative scores across 55 tasks from all benchmarks. The center line denotes the median. The box limits denote the first and third quartiles. The whiskers denote the full data range.

Pooling choice provides another view of the latent-space feature distribution. Within the evaluation framework of Section 2.2, we compared three widely used strategies, Classification Token pooling (CLS) [5, 18], Average pooling (AVG), and Max pooling (MAX). For the classification token strategy, we used the first token as the sequence representation for MLM-based and GenoJEPA architectures and the last token for NTP-based models. The distributions of fold-averaged task scores across 55 subtasks are shown as box plots in Figure 5b. Across the overall statistics, average pooling tended to give the strongest and most stable performance across most tasks [31]. Max pooling ranked second. This pattern suggests that aggregating features across the full sequence can better capture biological motifs that lack explicit boundaries. By contrast, the classification token strategy showed weaker overall performance, which indicates that a single token without task-specific finetuning may be insufficient to represent core contextual information in low signal-to-noise DNA sequences. GenoJEPA remained among the strongest models under all three tested pooling strategies.

To complement the pooling analysis, we further examined representation separability under different decision boundaries. We compared three classical machine learning classifiers, K-Nearest Neighbours (KNN) [39], Logistic Regression (LogReg) [30], and CatBoost [40], which represent distance-based, linear, and nonlinear tree-based decision spaces, respectively. All classifier hyperparameters followed the default settings of scikit-learn [33] and the CatBoost library. Each classification task was evaluated through 10-fold cross-validation with a held-out test set. The overall distribution of probe scores across 55 subtasks is summarized in Figure 5c. Among the tested foundation models, KNN generally yielded the lowest predictive performance. Notably, the nonlinear CatBoost did not show a consistent advantage over linear logistic regression and instead performed worse on several tasks. This trend was especially clear for the GenoJEPA models, which suggests that strong pretrained representations can support accurate classification with a simple linear model rather than with more complex nonlinear transformations that are more prone to overfitting. Consistent with Figure 5b, GenoJEPA remained competitive under each tested classifier.

Taken together, the few-shot, pooling, and classifier analyses support the quality of the native representations learned by GenoJEPA from multiple perspectives. The model generalized well with limited labelled data and achieved effective separation of complex biological functions in latent space. These aggregate summaries should still be interpreted alongside task-level heterogeneity, because some subtasks favoured particular baselines. Even with that caveat, the combination of data efficiency and representation separability remains attractive for practical biological research in settings where annotation and computation are both limited.

## 3 Discussion

Decoding the regulatory syntax embedded in genomic sequences remains a central challenge in computational biology. Most existing foundation models adopt sequence modelling paradigms derived from NLP [7, 19]. DNA sequences, however, lack explicit semantic boundaries and contain substantial neutral evolutionary noise. Precise nucleotide-level fitting in a low-dimensional discrete space may therefore encourage models to spend capacity on highfrequency details that are only weakly related to regulatory function [15]. To address this limitation, we introduced GenoJEPA, an unsupervised joint-embedding predictive architecture based on the LeJEPA formulation [23]. This design shifts learning from local reconstruction in input space to semantic alignment in a high-dimensional latent space. Across 55 downstream tasks, our results suggest that this design can learn transferable genomic representations with a favourable balance between performance and efficiency.

Another contribution of this work concerns the interaction between structural bias and computational efficiency in DNA tokenization. Conventional methods based on k-mer [7] or BPE [8, 19] are prone to vocabulary inflation and sensitive to single-nucleotide polymorphisms. Single-nucleotide tokenization [10, 22] avoids these problems, but it also introduces substantial computational overhead in the attention mechanism. Convolutional downsampling [24, 41] can reduce token length, yet it occupies a substantial fraction of the backbone parameter budget. The continuous patching strategy [18] used in GenoJEPA reduces the redundancy associated with discrete vocabularies while preserving biochemical dependencies in continuous space. Compared with single-nucleotide input, it shortens the effective sequence length without the overhead of stacked front-end computation.

The most practically important aspect of GenoJEPA is its ability to provide discriminative representations with frozen weights. This property is relevant for laboratories with limited computational resources that cannot routinely finetune large foundation models. Pretrained models that emphasize nucleotide reconstruction often struggle to yield equally strong frozen features [12]. By contrast, the probing results indicate that lightweight GPU-free classifiers such as logistic regression [30] can already achieve strong predictive performance from representations extracted by GenoJEPA across many tasks. This lowers the barrier to adopting genomic foundation models in biological laboratories. More broadly, the study suggests that latent-space semantic alignment is a useful direction for efficient training and broader application of larger-scale models in genomic research.

This work also has several limitations that motivate future research. The most immediate limitation concerns sequence length. Pretraining was conducted on sequences of up to 4,096 bp, which covers most gene-level tasks but is insufficient for analyses that require longer-range dependencies, such as topologically associating domain (TAD) boundary identification. The training advantages demonstrated in Section 2.4 suggest that GenoJEPA provides a practical basis for extending this limit. A further limitation concerns model scale. Although preliminary evidence for scaling laws [42] exists, the largest GenoJEPA model contains only 52M parameters, and the effects of further scaling remain unexplored.

## 4 Methods

### 4.1 Model architecture

#### 4.1.1 Tokenization

In genomic sequence modelling, tokenization determines how the model receives and represents biochemical signals. Current approaches mainly rely on BPE [8, 19] or k-mer tokenization [7, 20], both of which have clear limitations when applied to biological sequences. The frequent patterns captured by BPE do not necessarily correspond to functional conservation. The vocabulary size of k-mer tokenization grows exponentially with *k*, which makes large values computationally prohibitive. Both schemes may also fragment motifs that carry biological meaning and are sensitive to sequence variation. Even a single-nucleotide mutation can map a segment to a different identifier, obscure evolutionary relationships, and make it harder for the model to capture underlying biophysical and semantic similarity. Singlenucleotide tokenization [10, 22] avoids these issues, but it produces long token sequences and therefore incurs quadratic self-attention cost. Some studies [24, 41] use stacked convolutional layers for local downsampling, yet these layers consume a substantial share of the backbone parameter budget and increase memory overhead.

Given the similarity between genomic sequences and natural images in information density and structural organization, we adopted the continuous patching strategy [18] widely used in CV and applied it to DNA sequence processing. We first used a nucleotide-level tokenizer to convert the raw sequence into an index tensor **x** ∈ ℝ*^L^*, where *L* is the original DNA length. To ensure complete partitioning, we padded the tail of **x** with placeholder indices according to a preset patch size *P*, which was set to 16 in this study. This extended the length to the nearest integer multiple of *P* and yielded the padded tensor **x**_pad_ ∈ ℝ*^L^*, where *L*^′^ is the padded DNA length. We then segmented **x**_pad_ into a two-dimensional patch tensor **X**_patch_ ∈ ℝ^*N × P*^ using a sliding window of size *P*, where *N* = *L*^′^*/P* is the total number of patches. The stride was set equal to *P*, so partitioning was non-overlapping and maximized sequence compression.

After partitioning, the nucleotide units within each patch were mapped into a lowdimensional dense feature space to capture biochemical and physical similarities among bases at the input layer. Specifically, the discrete patch tensor **X**_patch_ was passed through a learnable embedding layer to produce a continuous feature tensor **E** ∈ ℝ^*N × P × D_embed_*^, where *D*_embed_ is the nucleotide embedding dimension and was set to 16 in this study. The local dense features **E** were then flattened along the spatial dimension and projected into a higher-dimensional latent space. This operation aggregated each patch of *P* nucleotides into a single continuous feature vector and yielded the tensor **H** ∈ ℝ^*N × D*_hidden_^ required by the backbone, where *D*_hidden_ is the hidden dimension of the Transformer encoder and was set to 384 or 512 in this study.

This transformation created a continuous input for the subsequent Transformer encoder. Compared with conventional k-mer tokenization, continuous patching with dense mapping reduces the parameter inflation associated with large discrete vocabularies. It shortens the effective sequence length to 1*/P* of the original length with relatively low computational cost. Achieving a similar compression ratio with conventional k-mers would require a vocabulary of about 4 billion entries. Unlike approaches that collapse long segments into discrete orthogonal identifiers, our strategy combines nucleotide-level representations with patch-level linear projection and preserves biochemical similarity within each segment in continuous space. A detailed analysis of the embedding strategy is provided in Appendix C.2.

#### 4.1.2 Backbone

The backbone of GenoJEPA is based on ModernBERT [25]. ModernBERT retains the advantages of bidirectional encoders for semantic extraction while incorporating several architectural advances from recent autoregressive LLMs [26–28]. This combination improves computational efficiency without sacrificing model quality on long sequences. In NLP, ModernBERT has shown strong accuracy and efficiency on tasks such as multi-document retrieval and ultra-long context modelling.

Relative to a vanilla Transformer [38], the backbone introduces several relevant changes. For positional encoding, it replaces conventional absolute position embeddings with Rotary Position Embedding (RoPE) [43]. RoPE injects relative position information through rotation in the complex domain and supports length extrapolation. For attention, the full ModernBERT design alternates global and local attention across layers to reduce cost while preserving long-range context. Global attention is applied every few layers to capture long-range dependencies, whereas the remaining layers use sliding-window local attention. Because the pretraining sequence lengths used here were relatively short, we did not enable this alternating attention mechanism. For regularization and optimization, the backbone uses pre-normalization, which places layer normalization before the multi-head attention and feed-forward layers and helps stabilize gradient propagation in deeper networks. The model also adopts a bias-free design by removing bias terms from most fully connected layers and layer normalization, which concentrates the parameter budget on the main feature transformation matrices.

On this basis, we defined two backbone variants to address different computational constraints. The lightweight version, GenoJEPA-T, has about 6M parameters. The base version, GenoJEPA-B, has about 52M parameters. GenoJEPA-T uses 4 full-attention layers with a hidden dimension of 384, an intermediate dimension of 768, and 12 attention heads. GenoJEPA-B uses 12 full-attention layers with a hidden dimension of 512, an intermediate dimension of 2,048, and 16 attention heads. Additional architectural details are provided in Table S2 in the Appendix.

#### 4.1.3 Optimization

Our goal was not to derive a new JEPA theory but to examine how this paradigm behaves in genomic representation learning. In self-supervised sequence modelling, MLM [5] and NTP [6] remain the dominant pretraining paradigms. Both natural images and genomic sequences, however, contain substantial redundancy and have relatively low signal-to-noise ratios because they arise through natural processes rather than deliberate design. Forcing models to reconstruct the input at high fidelity can therefore direct capacity towards low-dimensional and high-frequency local detail, which may limit abstraction of broader topological structure and high-dimensional semantics [15]. This often leads to weaker discrimination under probing without finetuning. JEPA [12] offers an alternative by aligning representations of views that share semantics in an abstract latent space. This avoids element-level reconstruction and encourages learning of the data manifold. Conventional JEPA implementations, however, often rely on heuristic devices such as asymmetric networks, momentum encoders, or stop-gradient operations to prevent representation collapse, where all inputs are mapped to nearly identical representations. These empirical designs increase hyperparameter sensitivity and may limit scalability and training stability. We therefore adopted the LeJEPA [23] framework, which avoids these heuristic interventions and derives feature regularization from a theoretically motivated objective. The inherited assumptions and their relevance to GenoJEPA are summarized in Appendix B.

For the optimization pipeline, we followed the general JEPA paradigm [17, 44]. Multiple views are produced by applying different augmentations to the same sample, and representations of matching views are brought closer together in latent space to learn invariances that are robust to local noise. We designed a DNA-specific multi-view augmentation strategy based on random cropping. For each sample, pretraining used 2 global views, with cropping scales uniformly sampled between 65% and 80% of the original sequence length, to capture global structure and long-range dependencies. It also used 6 local views, with cropping scales restricted to 35% to 40%, to emphasize local sequence variation and core biological motifs. Appendix C.1 complements this design with additional analyses of random mutation and reverse-complement augmentation. Each view was passed through the backbone and average pooling to obtain a view-level representation. A nonlinear projection layer with GELU activation then mapped this representation into a dedicated latent space of dimension 16 for loss computation. This module was active only during pretraining and helped isolate task-specific bias from the self-supervised objective, thereby preserving transferability of the backbone features [29, 45]. The pooling strategies and projection layer used during pretraining are analysed in Appendix C.2.

Within the dedicated latent space, GenoJEPA follows the LeJEPA [23] formulation rather than the I-JEPA [12] route. The representations of all views, both global and local, are aligned towards the mean representation of the global views. The model first computes the mean representation of all global views as the anchor target and then encourages all view representations to converge towards this anchor through Mean Squared Error (MSE). The invariance loss is defined as

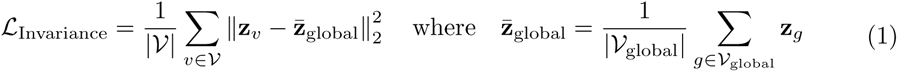

Here, 𝒱 and 𝒱_global_ denote the set of all views and the set of global views, respectively, and **z** represents the projected latent representation of each view.

Because this alignment operation tends to collapse the learned representations to a constant vector, a mechanism that preserves diversity is required. LeJEPA introduces SIGReg as a principled regularizer that steers latent features towards an isotropic Gaussian target without relying on additional empirical anti-collapse heuristics. SIGReg guides high-dimensional features towards this target by matching the Empirical Characteristic Function (ECF) to that of the standard Gaussian, given by 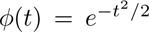. To enable efficient optimization, continuous integrals and expectations are approximated by discrete empirical estimates:

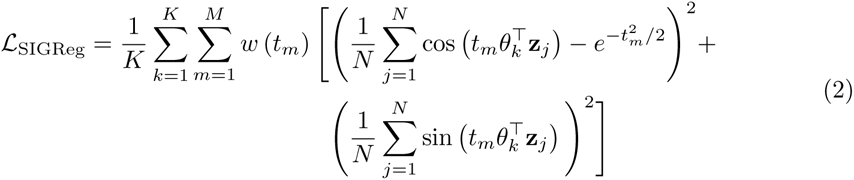

where *N* denotes the total number of representations **z***_j_* in a given training iteration, *θ_k_* is a unit direction vector drawn from the latent space, and *t_m_* and *w*(*t_m_*) represent the discrete evaluation points and their corresponding weights, respectively. In practice, the outer summation involves *K* = 256 random projection directions, where the unit direction vectors *θ_k_* are sampled from the latent space and the high-dimensional features **z***_j_* are projected onto the corresponding one-dimensional subspaces for evaluation. The inner summation performs numerical integration with the evaluation-domain upper bound *t_max_* set to 3.0. Evaluation points *t_m_* are uniformly sampled over [0, 3.0] to yield *M* = 17 discrete values, and integration weights *w*(*t_m_*) are applied to compute the discrete squared error.

Finally, to jointly optimize view alignment and representation distribution, network parameters are updated by minimizing a weighted combination of the invariance loss and the regularization term. The total loss is

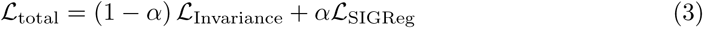

In this study, *α* was set to 0.05. In our experiments, this formulation yielded general-purpose representations with high linear separability in downstream tasks without requiring heuristic interventions. Sensitivity analyses for the loss hyperparameters are detailed in Appendix C.3.

### 4.2 Pretraining configuration

#### 4.2.1 Datasets

To train GenoJEPA, we used the multispecies genomic collection introduced in the Nucleotide Transformer study [7] as the pretraining corpus. This dataset was designed to balance reference-genome quality and taxonomic representativeness. A downsampling strategy was applied to the large number of bacterial taxa to avoid skew towards a single lineage. The resulting 850 representative species span major evolutionary branches, including bacteria, fungi, invertebrates, protozoa, and vertebrates. Using this diverse collection rather than a single species encouraged the model to capture evolutionary constraints and regulatory syntax across varied genomic contexts.

After obtaining the raw FASTA files of target species from NCBI [46], we applied a standardized sequence-cleaning and segmentation pipeline. All genomic sequences were first converted to uppercase, and non-canonical or ambiguous characters outside the standard A, T, C, G alphabet were replaced with N. To conform to the input requirements of the model, raw sequences were divided into fixed-length segments using a sliding-window strategy with a window size of 4,096 bp and an overlap of 512 bp between adjacent windows. This overlap helps preserve local sequence context and reduces the risk of truncating cis-regulatory elements or functional motifs at window boundaries. Fragments in which unknown nucleotides exceeded 15% of the total length were discarded.

After filtering and segmentation, the final dataset balanced representativeness and diversity across taxonomic groups. Mammalian vertebrates account for the largest share of the corpus, with 31 species contributing more than 21.3 million sequence fragments and about 87.1 billion effective nucleotides. Other vertebrates also contribute substantially, with 57 species producing more than 76.6 billion nucleotides. Invertebrates from 39 species contribute nearly 24.5 billion nucleotides and further increase structural diversity. Although individual microbial and fungal genomes are relatively compact, these groups dominate in species count. In total, 667 bacterial species and 46 fungal species contribute more than 4.7 billion nucleotides, and 10 protist species contribute more than 0.4 billion nucleotides. Overall, the corpus spans about 193 billion nucleotides. Detailed statistics for each taxonomic group are provided in Table S1 in the Appendix.

#### 4.2.2 Pretraining

Pretraining was conducted on the genomic corpus described above. The model was optimized with AdamW [47], with momentum parameters *β*_1_ and *β*_2_ set to 0.9 and 0.999, respectively. The numerical stability parameter *ɛ* was set to 1 × 10^−8^. We applied global gradient clipping with a maximum norm of 1.0 to maintain stable training on genomic data with low signalto-noise ratios. The learning rate increased linearly from 0 to 5 × 10^−4^ over the first 5,000 steps and then decayed with a cosine schedule to 5 × 10^−5^. Weight decay increased from 0.005 to 0.05 under a cosine schedule. This setup allowed broader parameter exploration early in training and stronger regularization later. Sensitivity analyses for learning rate and weight decay are provided in Appendix C.4. GenoJEPA-T, which has about 6M parameters, was trained on a single RTX 3090 24 GB GPU with a global batch size of 256 for 100,000 update steps and required about 12 hours. GenoJEPA-B, which has about 52M parameters, was trained on a single A800 80 GB GPU with the same batch size for 500,000 steps to better cover the larger pretraining corpus and required about 150 hours.

### 4.3 Evaluation protocol

#### 4.3.1 Benchmarks

The experiments covered three evaluation benchmarks, Genomic Benchmarks, GUE Benchmarks, and Nucleotide Transformer Tasks. Genomic Benchmarks and GUE Benchmarks include multispecies data, whereas Nucleotide Transformer Tasks focuses on the human genome. Together, these resources contain 55 subtasks that span cross-species sequence classification, promoter and enhancer detection, RNA splice site recognition, transcription factor binding prediction, and chromatin mark identification. For each task, we retained the benchmark-defined held-out test split and treated the remaining labelled data as a development pool divided into 10 folds. In each round, 9 folds were used for model fitting, 1 fold was used for early stopping, and the trained model was evaluated on the fixed held-out test set. For readability, we simplified some benchmark and task labels in the text and figures. The corresponding mappings are provided in Table S26 in the Appendix.

##### Genomic Benchmarks

The Genomic Benchmarks [21] provide a standardized and biologically informative framework for genomic sequence classification. It is assembled from public resources such as Ensembl together with experimental data from prior studies. The dataset includes sequences from model organisms such as human, mouse, and Caenorhabditis elegans, as well as multispecies datasets for comparative genomics. The collection contains 9 subtasks, including coding versus intergenic sequence classification and identification of regulatory elements such as enhancers, promoters, and open chromatin regions. Sequence lengths range from 2 bp to 4,776 bp, and sample sizes range from 968 to 231,348. This diversity requires models to capture genetic information across a wide range of sequence lengths and sample scales.

##### GUE Benchmarks

The GUE Benchmarks [19] extend evaluation to multispecies genomic language model understanding. Released with the DNABERT-2 study, it contains 28 subtasks covering data from human, mouse, yeast, and viral genomes, with a focus on transcriptional regulation and epigenetic assessment. The tasks include nine-class prediction of SARS-CoV-2 variants on sequences of 1,000 bp with more than 70,000 samples, binary classification of yeast histone modification states on sequences of 290 to 500 bp, and prediction of human and mouse transcription factor binding sites at 101 bp. Additional tasks include core and general promoter detection at 70 bp and 300 bp, and multi-class RNA splice site recognition at 400 bp. Sample sizes range from thousands to tens of thousands, allowing assessment across diverse biological signals.

##### Nucleotide Transformer Tasks

The Nucleotide Transformer Tasks [7] focus on functional element analysis in the human genome. Its 18 subtasks were developed by the Nucleotide Transformer team using data from ChIP-seq experiments in ENCODE [48] and the EPD database. For chromatin landscape analysis, models perform binary classification of 10 histone marks and epigenetic modifications on sequences of 1,000 bp. For regulatory element detection, the tasks include identifying TATA boxes in promoter regions with 300 bp sequences and detecting enhancers together with tissue specificity using 400 bp sequences. For RNA splicing, models classify splice donors and acceptors on 600 bp sequences. Each task typically provides up to 30,000 balanced training samples per class and therefore supports rigorous evaluation of transcriptional and epigenetic regulation in the human genome.

#### 4.3.2 Baselines

We selected five representative genomic foundation models as baselines, HyenaDNA, CaduceusPh, GROVER, DNABERT-2, and NT-v2. Together, these models cover the main tokenization strategies and backbone families used in the field. All baselines were initialized with publicly available optimal pretrained weights. Detailed model statistics are provided in Table S3 in the Appendix.

##### HyenaDNA

HyenaDNA [22] adopts a decoder-based architecture that replaces conventional attention with the Hyena operator [37], which combines long convolutions, implicit parameterization, and data-controlled gating to achieve sub-quadratic complexity. It tokenizes sequences at single-nucleotide resolution and is pretrained on a single human reference genome through autoregressive NTP. We used the LongSafari/hyenadna-large-1m-seqlen-hf version with 7M parameters and an output embedding dimension of 256.

##### CaduceusPh

CaduceusPh [10] applies selective State-Space Models (SSM) [49] to genomic modelling. Built on the Mamba [36] architecture, it achieves bidirectional sequence modelling through BiMamba modules and includes dedicated components to maintain reverse-complement equivariance. It tokenizes each sequence at single-nucleotide resolution and is pretrained on a single human reference genome using MLM. We used the kuleshovgroup/caduceus-ph seqlen-131k d model-256 n layer-16 version with 8M parameters and an output embedding dimension of 256.

##### GROVER

GROVER [8] is a 12-layer bidirectional Transformer [38] encoder designed to learn the intrinsic grammar of the human genome. It introduces BPE [50] for DNA tokenization, producing variable-length tokens that balance vocabulary frequency and reflect the heterogeneous composition of the genome. The model is pretrained on a single human reference genome using MLM. We used the PoetschLab/GROVER version with 87M parameters and an output embedding dimension of 768.

##### DNABERT-2

DNABERT-2 [19] adopts a BERT-like bidirectional Transformer encoder, replacing absolute position embeddings with Attention with Linear Biases (ALiBi) [51] to enhance length extrapolation. It uses a BPE tokenizer and is pretrained on the genomes of 135 species using MLM. We used the zhihan1996/DNABERT-2-117M version with 117M parameters and an output embedding dimension of 768.

##### NT-v2

NT-v2 [7] is a Transformer encoder that employs fixed 6-mer tokenization with MLM. It is pretrained on the genomes of 850 species and introduces RoPE [43] combined with a bias-free design to reduce parameter count and computational cost. We used the InstaDeepAI/nucleotide-transformer-v2-500m-multi-species version with 494M parameters and an output embedding dimension of 1,024.

#### 4.3.3 Metrics

Class imbalance is a common challenge in bioinformatics classification. Conventional metrics may not reflect true performance under skewed label distributions because they often neglect true negatives or favour the majority class. We therefore adopted MCC [34] as the primary evaluation metric throughout all classification tasks. MCC is a discrete form of the Pearson correlation coefficient between predicted and true labels. It incorporates all four elements of the confusion matrix and yields a high score only when the model performs well across all classes. MCC ranges from -1 to 1, where 1 indicates perfect prediction and 0 indicates random guessing.

We used the multiclass form of MCC implemented in scikit-learn [33], expressed as

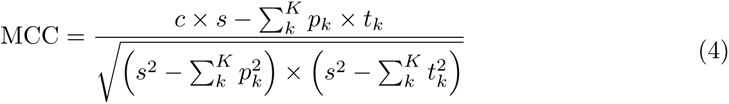

where *K* is the total number of classes, *c* is the sum of diagonal elements of the confusion matrix representing the total number of correctly classified samples, and *s* is the total number of samples. For class *k*, *p_k_* denotes the number of samples predicted as that class, and *t_k_* denotes the actual number of samples belonging to that class. The numerator captures the covariance between true and predicted distributions, while the denominator provides standard deviation normalization.

#### 4.3.4 Probing

To assess representation quality, all pretrained backbones were kept fully frozen during probing. Unless stated otherwise, we applied average pooling to the last-layer token representations and appended a logistic regression classifier [30]. Average pooling aggregates global information without introducing additional parameters. Logistic regression is computationally efficient and avoids confounds introduced by more complex fitting procedures. Downstream classifiers used default settings from standard machine learning libraries [33] to assess out-of-the-box performance. Because class imbalance is common across genomic tasks, MCC [34] was used consistently as the primary metric. To reduce dependence on any single train-validation split, each task was evaluated with 10-fold cross-validation with a held-out test set. For each task, the model was trained on 9 folds and validated on the remaining fold, yielding 10 models that were then evaluated on the test set. Final task performance is reported as the mean over these 10 test-set evaluations.

##### Main probing experiment

In the main probing experiment, we evaluated the semantic discriminability of each foundation model in latent space. Paired Wilcoxon signed-rank tests at *p* = 0.05 were applied to the 10 fold-wise test MCC values for each task to summarize paired performance under the common protocol. Because these fold-wise evaluations share the same held-out test split within each task, we treated them as a stability-aware comparison rather than as fully independent repeated samples. For overall presentation, we first computed the mean test score across folds for each task and then computed the mean and median across all tasks within each benchmark to construct radar plots. The same task-level means were used for subtask-level analysis and for the overall probing summaries across all benchmarks. To compare overall rankings, Friedman tests at *p* = 0.05 were performed on the task means, followed by Nemenyi post-hoc tests.

##### Few-shot probing experiment

To examine the generalization capacity of each model under data-scarce settings, we further conducted few-shot probing evaluations. The test set was kept unchanged throughout. Within each cross-validation round, we stratified the current fitting and validation subsets at proportions ranging from 10% to 50% within each class label. The same downsampled splits were used for all methods. Logistic regression classifiers and average pooling were applied consistently. For each task at each sampling proportion, the reported score is the mean test MCC across the 10 cross-validation rounds. The final reported performance was computed by aggregating these task-level means across all 55 tasks from the three benchmarks.

##### Pooling strategy analysis

To evaluate how feature aggregation affects sequence-level representation, we compared three pooling strategies. AVG averages all token embeddings except padding tokens to produce a global representation. MAX selects the maximum value along each feature dimension, again excluding padding tokens. The third strategy uses the CLS token [5]. For models trained with MLM or JEPA, namely DNABERT-2, NTv2, GROVER, GenoJEPA-T, and GenoJEPA-B, we extracted the first special token as the sequence representation. For models based on NTP, namely HyenaDNA and CaduceusPh, we extracted the last token to capture the full sequential context. Each strategy was evaluated over the same 10 cross-validation rounds for each task, and the distribution of task-level mean values across 55 tasks is summarized with box plots.

##### Classifier strategy analysis

To examine representation separability under different decision boundaries, we used three classifiers, KNN [39], logistic regression, and CatBoost [40], which cover distance-based, linear, and tree-based decision rules. KNN and logistic regression were implemented with scikit-learn [33], and CatBoost was implemented with its official library. All three models used the default settings of their respective frameworks. KNN used 5 neighbours with uniform weights and Euclidean distance. Logistic regression used L2 regularization with a penalty coefficient of 1.0 and a maximum of 100 iterations. CatBoost used a maximum tree depth of 6, an L2 regularization coefficient of 3, and a maximum of 500 iterations. Each classifier was evaluated over the same 10 cross-validation rounds for each task, and the distribution of task-level mean values across 55 tasks is summarized with box plots.

#### 4.3.5 Finetuning

We conducted full-parameter finetuning experiments to estimate the upper bound of recognition accuracy for each model. This setup allowed pretrained parameters to adapt fully to the distribution of each downstream task. We retained the model-specific pooling strategies and classification heads recommended in the original publications to approximate the strongest practical adaptation regime for each model, while probing remained strictly unified to support fair comparison of representations. Specifically, DNABERT-2, NT-v2, and GROVER used the first classification token embedding, HyenaDNA used the last non-padding token representation, and CaduceusPh used average pooling. GenoJEPA also used the first classification token embedding, followed by an MLP head composed of two linear layers with GELU [35] activation and a hidden dimension matched to the backbone. The effects of different pooling strategies during finetuning are discussed in Appendix C.5.

Core hyperparameters were searched over a grid of batch sizes {64, 128, 256} and learning rates {5 × 10^−5^, 1 × 10^−4^, 5 × 10^−4^, 1 × 10^−3^}. The best configurations for each model and dataset are reported in Table S27 in the Appendix. All models were optimized with AdamW [47], with *β*_1_ = 0.9, *β*_2_ = 0.999, and *ɛ* = 1 × 10^−8^. We also applied gradient clipping with a maximum norm of 1.0. Training ran for at most 100 epochs. The learning rate followed a cosine annealing schedule with linear warmup over the first 10% of training and then decayed to 10% of the peak value. Weight decay followed a cosine schedule and increased from 0.001 to 0.01 over training. MCC was used as the primary metric. Early stopping was applied to balance convergence and overfitting. Training stopped and weights were rolled back to the best checkpoint when validation MCC failed to improve for 10 consecutive evaluation cycles. All finetuning experiments were conducted on a single A800 80 GB GPU.

##### Main finetuning experiment

For each of the 55 subtasks, finetuning was evaluated with the same 10-fold cross-validation with held-out test set used in probing. The mean MCC across the 10 fold-wise test evaluations was used as the representative score for each subtask. This task-level mean was used consistently for metric aggregation, both for subtask-level reporting and for overall summaries across all benchmarks. To compare model rankings, Friedman tests at *p* = 0.05 were performed on the task-level means, followed by Nemenyi post-hoc tests.

##### Computational efficiency analysis

To evaluate the deployment potential of different backbones, we measured runtime and memory usage. All tests were executed on a single A800 80 GB GPU with a batch size of 1, following the protocol in [31], and covered sequence lengths from short fragments to beyond the hardware limit. To reduce noise from system scheduling and obtain stable estimates, each test was run independently for 10 epochs. Within each epoch, the first 10 steps were treated as warmup and excluded. The mean of the next 100 steps was used to represent core performance. The final metric at each sequence length was the mean of these 10 epoch-level estimates.

## Data availability

The 850 species used for pretraining are listed at https://huggingface.co/datasets/InstaDeepAI/multi_species_genomes. The corresponding FASTA files can be downloaded from NCBI at https://ftp.ncbi.nlm.nih.gov/genomes/refseq. Given the large file sizes, we provide only the preprocessing code, as described in the Code availability section. The raw Genomic Benchmarks, GUE Benchmarks, and Nucleotide Transformer Tasks datasets are available at https://huggingface.co/katarinagresova/datasets, https://github.com/MAGICS-LAB/DNABERT_2, and https://huggingface.co/datasets/InstaDeepAI/nucleotide_transformer_downstream_tasks_revised, respectively. The preprocessed evaluation datasets will be made publicly available upon publication.

## Code availability

All preprocessing code, training and evaluation code, and preprocessed datasets will be made publicly available upon publication.

## Acknowledgements

This work was supported in part by the National Key R&D Program of China 2024YFE0200800, the National Natural Science Foundation of China under Grants U23B2001 (62471055, 62321001, 62401080, 62101064, 62171057, 62201072, 62071067).

## Author contributions

C.W. and J.W. conceived and designed the study. C.W. designed the model architecture and led the pretraining pipeline. Q.Q. designed the evaluation experiments and the dataprocessing pipeline. B.H. collected and curated the datasets. S.L. conducted the evaluation experiments. H.S. advised on data analysis and interpretation. Z.Z. performed the theoretical analysis. J.W. and J.L. provided computational support and analysis tools. C.W. and Q.Q. wrote the manuscript with input from all co-authors.

## A Related work

### A.1 Representation learning paradigms

The growing availability of unlabelled data has gradually reduced the annotation bottleneck in supervised learning and established self-supervised representation learning as a central paradigm. In Natural Language Processing (NLP), Masked Language Modelling (MLM) as exemplified by BERT [5] and Next-Token Prediction (NTP) as represented by GPT [6] have led to major advances. Because text tokens are already highly compressed abstractions of human thought, reconstructing or predicting them forces models to perform deep syntactic parsing and broad contextual reasoning. Inspired by this success, Computer Vision (CV) introduced Masked Autoencoders (MAE) [14], which reconstruct masked image patches in pixel space. Images, however, contain substantial redundant detail and have not undergone semantic compression. As a result, pixel-level reconstruction can bias networks towards local texture and other low-level features. Although MAE provides effective initialization for finetuning, the resulting features often show limited semantic discrimination when probed with a frozen backbone [15].

To overcome this bias, joint-embedding predictive architectures (JEPA) shift the learning target to a high-dimensional latent space. Early contrastive methods such as SimCLR [44] and MoCo [45], together with non-contrastive methods such as BYOL [29] and DINO [17], align different augmented views of the same image and suppress irrelevant noise. More recent approaches such as I-JEPA [12] and V-JEPA [52] predict target-view representations directly in latent space, which filters fine-grained detail and yields semantically richer features. Despite their effectiveness, conventional implementations still rely on momentum teacher networks or stop-gradient operations to prevent representation collapse. These mechanisms lack rigorous mathematical guarantees and introduce hyperparameter sensitivity. The LeJEPA [23] framework addresses this limitation through Sketched Isotropic Gaussian Regularization (SIGReg), which guides latent features towards an isotropic Gaussian distribution that is theoretically motivated to generalize well across unknown downstream tasks. By replacing heuristic designs with this theoretically grounded objective, LeJEPA reduces tuning cost and improves both discriminability and training stability.

### A.2 Genomic foundation models

In computational biology, decoding the regulatory syntax embedded in non-coding DNA remains a central challenge. Work in this area has progressed along two main directions. The first relies on supervised learning to map raw DNA sequences directly to functional genomics profiles. For example, Enformer [24] combined convolutional and Transformer modules to predict gene expression and chromatin states, and AlphaGenome [41] extended the context window to the megabase scale. However, high-quality experimental annotations remain limited, and supervised models are often tied to specific pretraining tasks or biological contexts.

Because unlabelled nucleotide data are far more abundant than annotated data, selfsupervised genomic language models have emerged as a complementary direction. Models such as DNABERT [9, 19], GROVER [8], Nucleotide Transformer [7], and CaduceusPh [10] adapted MLM to capture local biological motifs and cis-regulatory interactions. HyenaDNA [22] and GENERator [20] used NTP for autoregressive pretraining and highlighted the potential to model longer-range dependencies. Architectures borrowed from NLP, including Transformers [38] with RoPE [43] and State-Space Models (SSM) [49] such as Mamba [36] and Hyena [37], have also been applied to DNA modelling. Despite clear progress, these models still operate mainly at the nucleotide level. As in MAE [14] for vision, fitting high-frequency noise at this level can bias representations towards low-level physical detail rather than higher-level semantics, and downstream adaptation often still requires substantial finetuning [15]. The present study therefore introduces JEPA [23] to genomic sequence modelling, uses latent-space semantic alignment to attenuate nucleotide-level noise, and aims to provide an off-the-shelf feature extractor that can be used without parameter updates.

Beyond the pretraining objective, the conversion of nucleotides into neural network inputs is equally important. Existing tokenization strategies, including Byte-Pair Encoding (BPE) [8, 19], fixed-window k-mer segmentation [7, 20], and single-nucleotide tokenization [10, 22], each have clear limitations. The statistical patterns learned by BPE do not necessarily correspond to functionally meaningful conservation. The vocabulary size of k-mer tokenization grows exponentially with *k*, which makes large values impractical. Both schemes may fragment biologically meaningful motifs and are highly sensitive to sequence variation, because even a single mutation can map a segment to a different token. Single-nucleotide tokenization avoids these issues but creates long token sequences and therefore incurs quadratic self-attention cost. Some studies [24, 41] have introduced convolutional downsampling, yet stacked convolutional layers occupy a substantial part of the parameter budget. In this work, we instead adopt continuous patching with linear projection [18], which avoids discrete vocabulary inflation while preserving biochemical dependencies within each segment.

## B Theoretical formulation

This section summarizes the theoretical framework inherited from LeJEPA [23] and explains how it motivates the design choices adopted in GenoJEPA. The aim is to clarify the rationale for the pretraining objective rather than to claim new proofs beyond the original LeJEPA analysis.

### B.1 Isotropic Gaussian optimality

In the LeJEPA analysis, the theoretical goal of JEPA-style representation learning is to learn general representations that transfer well to unseen downstream tasks. To formalize which feature distribution is optimal, the analysis minimizes prediction risk without assuming a specific downstream task. Consider first linear probing. When downstream decision boundaries are obtained through ordinary least squares with Tikhonov regularization [53], the theory shows that anisotropic feature covariance, where variance is unevenly distributed across dimensions, amplifies the statistical bias of the linear estimator in a nonlinear manner. Even in the high-variance regime where regularization is removed, the trace of the estimator variance remains strictly larger for anisotropic features than for isotropic ones. An isotropy constraint is therefore necessary for low-bias and low-variance predictions.

When the analysis is extended to nonlinear probes such as radius-based K-Nearest Neighbours [54] and Nadaraya-Watson kernel regression [55], the necessity and uniqueness of the isotropic Gaussian distribution can be established through variational calculus. For nonlinear estimators, generalization error is dominated by the integrated squared bias of the density rather than by covariance alone. Through Taylor expansion and expectation analysis on local feature manifolds, this bias term is shown to be closely related to the Fisher information functional of the embedding density. Variational calculus together with the matrix Cramér-Rao lower bound shows that among all multivariate probability densities with a fixed total variance constraint, such as a trace or Frobenius norm constraint, the isotropic Gaussian distribution minimizes the Fisher information functional. In this setting, the Fisher information functional coincides with the integrated squared bias. This conclusion motivates treating representation collapse in self-supervised learning as a mathematical problem of distribution alignment.

### B.2 SIGReg objective formulation

Given this theoretical motivation for isotropic Gaussian features, the next challenge in LeJEPA is to design a differentiable and computationally efficient loss that drives highdimensional features towards the target distribution. Statistically, this is equivalent to performing a high-dimensional multivariate hypothesis test during each backward pass. Because the feature manifolds of modern neural networks often span hundreds of dimensions, direct computation of divergences or distances between multivariate joint distributions faces a severe curse of dimensionality. Sparse samples in high-dimensional space yield unstable gradients, and computational cost grows exponentially with dimension. Both issues are problematic in large-scale pretraining [56].

To overcome this computational bottleneck, LeJEPA introduces a sketching approach drawn from integral geometry and high-dimensional statistics. Its mathematical basis is the hyperspherical Cramér-Wold theorem [57], which states that two high-dimensional random vectors have identical joint distributions if and only if their one-dimensional projections onto all unit directions are also identically distributed. Based on this equivalence, SIGReg avoids direct distribution fitting in the full latent space. During each forward pass, the algorithm samples *K* unit direction vectors *θ_k_* in the latent space. Each feature **z***_j_* in the current batch is projected along each sampled direction into a scalar 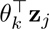. The discrepancy between this projected data distribution and the standard univariate Gaussian is then evaluated independently in each one-dimensional subspace. Reducing high-dimensional matching to a collection of one-dimensional comparisons lowers computational and memory cost to a scale that is linear in batch size. The method also exploits the implicit Sobolev smoothness [58] of deep network mappings, so that a relatively small number of projection directions can still constrain the full high-dimensional distribution effectively.

### B.3 Characteristic function test

After reducing distribution matching to one-dimensional projection spaces, the main question becomes which statistical test should be used to quantify the discrepancy between projected features and the target distribution, because this choice determines gradient stability during pretraining. Traditional moment-matching methods are intuitive, but they involve high-degree polynomial expansions when higher-order moments are computed. During backpropagation, gradient norms can therefore grow rapidly as moment order increases. In addition, matching only a finite number of moments cannot uniquely determine a continuous distribution and may allow shortcut solutions that circumvent the regularizer. Tests based on the Cumulative Distribution Function (CDF), such as the Cramér-von Mises test [59], present a different problem because they require exact sorting of all samples in the batch. Since sorting is non-differentiable, these tests require soft relaxations and also reduce synchronization efficiency in large-scale distributed training.

Following this comparison, SIGReg adopts the Epps-Pulley test [60], a frequency-domain test based on the Empirical Characteristic Function (ECF). In probability theory, the characteristic function is the Fourier transform of the probability density. For *N* one-dimensional projected features 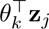 in the current batch, the ECF is defined as the arithmetic mean of the complex exponentials 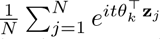, where *t* is the frequency-domain evaluation parameter. The standard univariate Gaussian used as the target distribution has a closed-form characteristic function that is purely real and given by 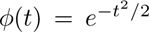.

To measure the discrepancy between the two, we compute the squared error integral in a weighted *L*_2_ space. By Euler’s formula *e^ix^* = cos *x* + *i* sin *x*, the complex-valued ECF can be decomposed into a real cosine component and an imaginary sine component. Because the characteristic function of the target Gaussian is purely real, with zero imaginary part, the squared complex distance at any frequency point *t* reduces to the sum of the squared real-part difference and the squared imaginary part. The continuous integral is then approximated with *M* discrete evaluation points *t_m_* and integration weights *w*(*t_m_*), which yields a discrete Riemann sum suitable for efficient end-to-end optimization. Aggregating this approximation across all *K* random projection directions gives the SIGReg loss in Equation 2.

An important property of this formulation is that sine and cosine are naturally bounded in [-1, 1]. As a result, the first-order gradients and second-order curvature of the loss are globally bounded by constants regardless of how extreme the input feature distribution becomes. This property helps prevent gradient explosion and contributes to smooth and stable large-scale pretraining.

### B.4 Unified pretraining objective

With this anti-collapse frequency-domain regularizer in place, integrating it with the semantic alignment mechanism of JEPA yields a simple pretraining objective under the inherited mathematical guarantees. In the standard self-supervised multi-view pipeline [17], input data are augmented and divided into global views that capture broad context and local views that focus on finer detail. The backbone extracts features, which are then mapped into a dedicated latent space through a nonlinear projection layer for loss computation.

To learn high-dimensional semantic invariances while suppressing low-level and highfrequency noise, the model aligns the representations of different views in latent space. Specifically, it computes the mean latent representation of the global views as an anchor semantic center and constrains all views to converge towards this center through mean squared error. This alignment defines the primary invariance loss 𝓛*_Invariance_*. On its own, however, this objective may allow a trivial solution in which all outputs collapse to a constant vector. The training loss therefore combines two objectives as in Equation 3, with a weighting hyperparameter *α* that balances semantic alignment and feature regularity.

Because SIGReg regularizes the feature distribution towards an isotropic Gaussian distribution, representational collapse is mitigated at the manifold level in LeJEPA [23]. As a result, the training paradigm does not require auxiliary components that are common in earlier self-supervised methods. During pretraining, the network does not rely on stop-gradient operations, momentum encoders, asymmetric predictor heads, or similar heuristic devices. This inherited argument motivates the GenoJEPA pretraining objective and is consistent with the separable representations observed in our downstream experiments.

## C Additional experiments

Pretraining genomic foundation models involves many hyperparameters and non-trivial architectural choices. To better understand how the model behaves on nucleotide sequences with low signal-to-noise ratios, this section reports 23 ablation analyses. Owing to computational constraints, all ablation studies used the lightweight GenoJEPA-T model. This version shares the same mathematical formulation as the larger models in terms of continuous patching, backbone architecture, and joint-embedding objective. Scaling mainly changes hidden-layer capacity and does not alter the core learning dynamics. The observed sensitivities to data augmentation, parameter settings, and architectural components therefore provide a useful reference for larger-scale models [61].

In architectural design and parameter selection, we followed a pragmatic trade-off. When increasing a parameter produced only marginal gains at a clear computational cost, we favoured a Pareto-optimal balance between performance and efficiency rather than the absolute peak score. Dataset splits and experimental settings were kept consistent with the main text. Unless noted otherwise, results for individual tasks are reported on the heldout test set and averaged over 10-fold cross-validation. Benchmark-level performance is the macro-average of task means within a benchmark, and cross-benchmark global metrics are computed across all 55 subtasks.

### C.1 Data augmentation

Species diversity in the pretraining corpus is important for model generalization. Table S4 compares training on a single human reference genome with training on a multispecies mixed corpus under the same number of iterations. A single human reference genome offers higher assembly quality, but it also contains substantial species-specific variation. By contrast, a multispecies corpus encourages the model to suppress species-specific noise in latent space and learn conserved cross-species patterns, which improves downstream generalization overall.

In JEPA, augmentation not only determines sample diversity but also defines the semantic invariances learned by the network. We primarily used multi-view cropping. As shown in Table S6, a moderate increase in the number of global views helps build a more stable semantic reference center. Excessive numbers of global views, however, produce diminishing gains, increase the memory burden, and may weaken learning of local motif features. Table S7 shows that moderately expanding the cropping range of the global views improves perception of broader topological structure. As the scale approaches the full sequence length, however, gains plateau while computational cost continues to rise.

Local view settings determine sensitivity to microscopic variation and fine-grained biological syntax. As shown in Table S8, increasing the number of local views improves the discriminability of high-dimensional features, although the marginal benefit decreases as computational cost rises. Table S9 shows that when local cropping roughly matches the spatial span of core regulatory elements, the model can align short-range variation with broader regulatory logic in latent space and improve feature resolution. As with the global views, further expansion of the local cropping scale yields diminishing returns while computational cost keeps increasing.

Beyond multi-view cropping, we also explored DNA-specific augmentation. Table S10 shows that increasing the random mutation probability progressively disrupts core regulatory motifs and generally reduces performance. Table S11 shows that increasing the reversecomplement probability improves performance on strand-symmetric tasks but impairs promoter and splice site recognition, which depends on transcriptional directionality. Taken together, these results suggest that unmodified sequence views provide the most balanced performance across tasks.

### C.2 Architecture design

The continuous patching strategy and low-level embedding mechanism are central to balancing long-sequence modelling against computational cost. Table S12 shows that smaller patch sizes provide finer local feature extraction but increase computational cost substantially. Excessively large patches, by contrast, may obscure critical mutation sites. A moderate patch size aligns well with the typical length of many regulatory elements. Table S13 shows that non-overlapping partitioning with stride equal to patch size reduces computational load markedly without an appreciable loss of accuracy. Table S5 shows that learnable continuous embeddings outperform one-hot input, presumably because they allow the model to capture underlying physicochemical properties of bases. Because genomic sequences contain only four bases and therefore limited information entropy, representational capacity saturates quickly as embedding dimension increases, as shown in Table S14. Compact low-dimensional embeddings are therefore sufficient for effective biochemical signal transmission.

We also examined the pretraining-specific modules placed between the backbone output and the loss computation. These modules help isolate pretraining-task bias and preserve backbone generalizability. Table S15 shows that average pooling performs best during pretraining, likely because it preserves global and continuous physical associations in an unbiased way. Table S17 shows that a nonlinear projection head is important for preserving the generalization of backbone features. When the alignment loss is applied directly to the backbone output, the deep weights become more tightly coupled to the specific cropping and alignment task. The projection layer acts as a semantic buffer and preserves transferability. Table S16 shows that batch normalization slightly outperforms layer normalization in the nonlinear projection head, possibly because it suppresses inter-species background variation and better matches the behaviour of SIGReg.

As shown in Table S18, a larger projection-head output dimension does not consistently improve performance. This behaviour reflects the way SIGReg approximates the distribution by randomly projecting high-dimensional features into one-dimensional subspaces. When the dimension is too high, approximation variance increases under a finite number of slices. When it is too low, an information bottleneck may appear. A moderate dimension therefore provides a practical balance between representational capacity and numerical stability.

### C.3 Objective function

The objective function not only drives parameter updates but also shapes the representation distribution in latent space. The distance criterion used for view alignment reflects the geometric assumptions imposed on that space. The comparison in Table S19 shows that cosine similarity ignores absolute feature magnitude and therefore fails to build clusters with clear boundaries. The linear penalty of absolute error can lead to gradient oscillation near the optimum. Mean squared error is better matched to Gaussian-distributed features. It places a stronger penalty on outlier views that deviate from the semantic center, promotes convergence of core motif vectors, and yields the best discriminability.

For SIGReg hyperparameters, Tables S20 and S21 show that moderate increases in the number of random projection slices and in the density of evaluation points improve coverage of the high-dimensional space. This reduces numerical approximation error in the ECF and guides the feature manifold more reliably towards the target Gaussian hypersphere. Excessive increases, however, yield diminishing returns while increasing computational cost. Table S22 shows that a moderate expansion of the integration domain helps capture tail behaviour, whereas an excessive expansion dilutes numerical weight in the core region and weakens the gradient signal.

The sensitivity analysis of the regularization weight in Table S23 shows the balance between alignment and diversity in feature space. When the weight is too low, the alignment objective dominates and features move towards local collapse. When it is too high, excessive repulsion disperses the features and disrupts alignment of homologous segments. A moderate weight produces a more uniform latent distribution while still allowing biochemically similar motifs to form compact local clusters.

### C.4 Optimization settings

When performing self-supervised pretraining on large multispecies genomic data, the joint strategy for learning rate and weight decay is critical for generalization. Compared with static settings, dynamic schedules allow the optimizer to adapt more flexibly across training stages. Table S24 shows that excessively high learning rates can lead to divergence and unstable gradients, whereas excessively low learning rates restrict the model from escaping local optima. A moderate initial learning rate combined with cosine decay supports broad exploration early in training and more refined feature learning later. Table S25 shows that excessive weight decay over-constrains the parameter space and impedes learning of important features, whereas insufficient regularization leaves the weights too unconstrained and promotes overfitting. A moderate weight-decay schedule with cosine warmup provides enough flexibility early in training and stronger regularization later.

### C.5 Downstream adaptation

During finetuning for specific downstream tasks, pooling strategy remains an important factor because it determines how semantic information from deep representations is transferred to the task head. Figure S1 shows behaviour that differs from what we observed during pretraining and probing. During pretraining, average pooling provides more comprehensive constraints on the representation space. During probing, the classification token is not task-specifically adapted and therefore cannot reliably aggregate information or suppress noise, whereas average pooling retains most information in a relatively unbiased way and is better suited to a lightweight downstream classifier. During finetuning, however, classification-token pooling shows strong overall performance and stable behaviour across all three benchmarks. This result is consistent with the objective of finetuning, which is to map general-purpose features to task-specific decision boundaries. In this setting, the classification token is optimized jointly with the classifier and gradually learns to aggregate task-relevant regulatory signals while reducing redundant background context that average pooling may retain. Max pooling, by contrast, tends to amplify non-specific repetitive signals.

---

**Fig. S1.**
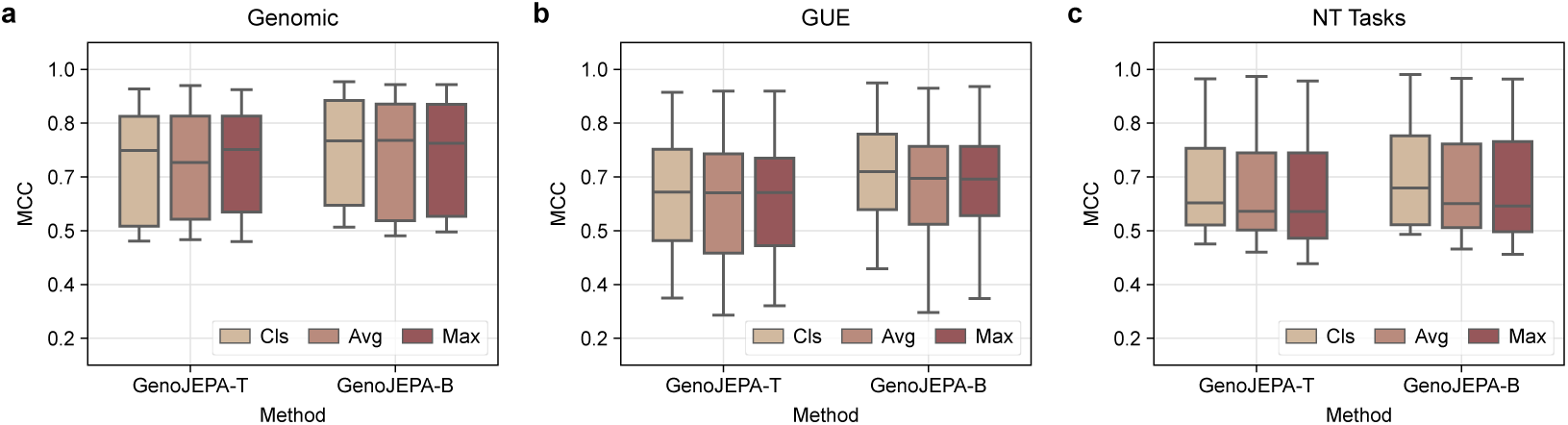
Finetuning performance of GenoJEPA with different pooling strategies for each benchmark. Each task is evaluated through 10-fold cross-validation, and the mean test score is used as the representative result. **(a)**, **(b)**, and **(c)** show results on the Genomic Benchmarks, the GUE Benchmarks, and the Nucleotide Transformer Tasks. Boxplots summarize the distribution of these mean scores across tasks within each benchmark. The center line denotes the median. The box limits denote the first and third quartiles. The whiskers denote the full data range.

**Table S1.** Summary statistics of the pretraining dataset. *Processed Sequences* and *Processed Nucleotides* refer to the statistics of segmented sequences.

| Taxonomic Group | Species Count | Raw Nucleotides | Processed Sequences | Processed Nucleotides |
| --- | --- | --- | --- | --- |
| Bacteria | 667 | 2,792,859,114 | 757,734 | 3,103,678,464 |
| Fungi | 46 | 1,524,710,503 | 400,899 | 1,642,082,304 |
| Invertebrate | 39 | 22,909,079,779 | 5,991,259 | 24,540,196,864 |
| Protozoa | 10 | 418,279,679 | 106,176 | 434,896,896 |
| Vertebrate (Mammalian) | 31 | 79,117,166,685 | 21,260,858 | 87,084,474,368 |
| Vertebrate (Other) | 57 | 71,657,061,222 | 18,705,801 | 76,618,960,896 |

**Table S2.** Architectural hyperparameters of GenoJEPA at different scales.

| Parameter | GenoJEPA-T | GenoJEPA-B |
| --- | --- | --- |
| Vocabulary Size | 5 | 5 |
| Embedding Size | 16 | 16 |
| Patch Size | 16 | 16 |
| Hidden Size | 384 | 512 |
| Intermediate Size | 768 | 2048 |
| Hidden Layers | 4 | 12 |
| Attention Heads | 12 | 16 |

**Table S3.** Architectural configurations of the baseline models.

| Model | Parameters | Architecture | Tokenizer | Embedding Dimension | Pretraining Task | Pretraining Species |
| --- | --- | --- | --- | --- | --- | --- |
| HyenaDNA | 7M | Hyena | Single Nucleotide | 256 | NTP | Human |
| CaduceusPh | 8M | Mamba | Single Nucleotide | 256 | MLM | Human |
| DNABERT-2 | 117M | Transformer | BPE | 768 | MLM | Multispecies |
| GROVER | 87M | Transformer | BPE | 768 | MLM | Human |
| NT-v2 | 494M | Transformer | 6-mer | 1024 | MLM | Multispecies |
| GenoJEPA-T | 6M | Transformer | Single Nucleotide | 384 | JEPA | Multispecies |
| GenoJEPA-B | 52M | Transformer | Single Nucleotide | 512 | JEPA | Multispecies |

**Table S4.** Performance of GenoJEPA with different genomic corpora across benchmarks. Each task is evaluated through 10-fold cross-validation, and the mean test score is used as the representative result. Average performance across tasks within each benchmark is reported from these mean scores. *Avg Score* denotes the overall average across 55 tasks from three benchmarks. The optimal results are marked in bold.

| Data Source | Genomic Benchmarks |  | GUE Benchmarks |  | NT Tasks |  | Avg. Score |  |
| --- | --- | --- | --- | --- | --- | --- | --- | --- |
|  | Finetuning | Probing | Finetuning | Probing | Finetuning | Probing | Finetuning | Probing |
| Human | 0.7082 | 0.5838 | 0.5822 | 0.5010 | 0.6549 | **0.5353** | 0.6266 | 0.5258 |
| Multispecies | **0.7140** | **0.5886** | **0.6278** | **0.5336** | **0.6656** | 0.5263 | **0.6543** | **0.5402** |

**Table S5.** Performance of GenoJEPA with different embedding strategies across benchmarks. Each task is evaluated through 10-fold cross-validation, and the mean test score is used as the representative result. Average performance across tasks within each benchmark is reported from these mean scores. *Avg Score* denotes the overall average across 55 tasks from three benchmarks. The optimal results are marked in bold.

| Embedding Strategy | Genomic Benchmarks |  | GUE Benchmarks |  | NT Tasks |  | Avg. Score |  |
| --- | --- | --- | --- | --- | --- | --- | --- | --- |
|  | Finetuning | Probing | Finetuning | Probing | Finetuning | Probing | Finetuning | Probing |
| One Hot | **0.7313** | **0.5905** | 0.6169 | 0.5213 | 0.6551 | 0.5146 | 0.6481 | 0.5304 |
| Embedding | 0.7140 | 0.5886 | **0.6278** | **0.5336** | **0.6656** | **0.5263** | **0.6543** | **0.5402** |

**Table S6.** Performance of GenoJEPA with different global view numbers across benchmarks. Each task is evaluated through 10-fold cross-validation, and the mean test score is used as the representative result. Average performance across tasks within each benchmark is reported from these mean scores. *Avg Score* denotes the overall average across 55 tasks from three benchmarks. The optimal results are marked in bold.

| Global Number | Genomic Benchmarks |  | GUE Benchmarks |  | NT Tasks |  | Avg. Score |  |
| --- | --- | --- | --- | --- | --- | --- | --- | --- |
|  | Finetuning | Probing | Finetuning | Probing | Finetuning | Probing | Finetuning | Probing |
| 1 | 0.7042 | 0.5915 | 0.6143 | 0.5246 | 0.6585 | 0.5200 | 0.6435 | 0.5340 |
| 2 | **0.7140** | 0.5886 | 0.6278 | 0.5336 | **0.6656** | 0.5263 | 0.6543 | 0.5402 |
| 4 | 0.7057 | **0.5971** | **0.6361** | **0.5338** | 0.6632 | **0.5292** | **0.6564** | **0.5427** |

**Table S7.** Performance of GenoJEPA with different global view scales across benchmarks. Each task is evaluated through 10-fold cross-validation, and the mean test score is used as the representative result. Average performance across tasks within each benchmark is reported from these mean scores. *Avg Score* denotes the overall average across 55 tasks from three benchmarks. The optimal results are marked in bold.

| Global Scale | Genomic Benchmarks |  | GUE Benchmarks |  | NT Tasks |  | Avg. Score |  |
| --- | --- | --- | --- | --- | --- | --- | --- | --- |
|  | Finetuning | Probing | Finetuning | Probing | Finetuning | Probing | Finetuning | Probing |
| [0.60, 0.75] | 0.6924 | 0.5788 | 0.6011 | 0.5126 | 0.6436 | 0.5129 | 0.6299 | 0.5235 |
| [0.65, 0.80] | 0.7140 | 0.5886 | 0.6278 | 0.5336 | 0.6656 | **0.5263** | 0.6543 | 0.5402 |
| [0.70, 0.85] | **0.7231** | 0.5911 | 0.6256 | 0.5317 | 0.6634 | 0.5256 | 0.6539 | 0.5394 |
| [0.75, 0.90] | 0.7154 | **0.6013** | 0.6294 | 0.5276 | 0.6643 | 0.5238 | 0.6549 | 0.5384 |
| [0.80, 0.95] | 0.7184 | 0.5937 | 0.6277 | **0.5352** | **0.6678** | 0.5261 | 0.6557 | **0.5418** |
| [0.85, 1.00] | 0.7179 | 0.5986 | **0.6355** | 0.5298 | 0.6607 | 0.5258 | **0.6572** | 0.5398 |

**Table S8.** Performance of GenoJEPA with different local view numbers across benchmarks. Each task is evaluated through 10-fold cross-validation, and the mean test score is used as the representative result. Average performance across tasks within each benchmark is reported from these mean scores. *Avg Score* denotes the overall average across 55 tasks from three benchmarks. The optimal results are marked in bold.

| Local Number | Genomic Benchmarks |  | GUE Benchmarks |  | NT Tasks |  | Avg. Score |  |
| --- | --- | --- | --- | --- | --- | --- | --- | --- |
|  | Finetuning | Probing | Finetuning | Probing | Finetuning | Probing | Finetuning | Probing |
| 2 | 0.6948 | 0.5876 | 0.6065 | 0.5282 | 0.6535 | **0.5292** | 0.6363 | 0.5382 |
| 6 | **0.7140** | 0.5886 | 0.6278 | 0.5336 | 0.6656 | 0.5263 | 0.6543 | 0.5402 |
| 14 | 0.7113 | **0.5988** | **0.6310** | **0.5396** | **0.6705** | 0.5291 | **0.6571** | **0.5459** |

**Table S9.** Performance of GenoJEPA with different local view scales across benchmarks. Each task is evaluated through 10-fold cross-validation, and the mean test score is used as the representative result. Average performance across tasks within each benchmark is reported from these mean scores. *Avg Score* denotes the overall average across 55 tasks from three benchmarks. The optimal results are marked in bold.

| Local Scale | Genomic Benchmarks |  | GUE Benchmarks |  | NT Tasks |  | Avg. Score |  |
| --- | --- | --- | --- | --- | --- | --- | --- | --- |
|  | Finetuning | Probing | Finetuning | Probing | Finetuning | Probing | Finetuning | Probing |
| [0.20, 0.25] | 0.7048 | 0.5785 | 0.6099 | 0.5096 | 0.6493 | 0.5061 | 0.6383 | 0.5197 |
| [0.25, 0.30] | 0.6968 | 0.5889 | 0.6216 | 0.5203 | 0.6561 | 0.5061 | 0.6452 | 0.5269 |
| [0.30, 0.35] | 0.7020 | **0.5939** | 0.6254 | 0.5244 | 0.6566 | 0.5111 | 0.6481 | 0.5314 |
| [0.35, 0.40] | **0.7140** | 0.5886 | **0.6278** | **0.5336** | **0.6656** | **0.5263** | **0.6543** | **0.5402** |

**Table S10.**
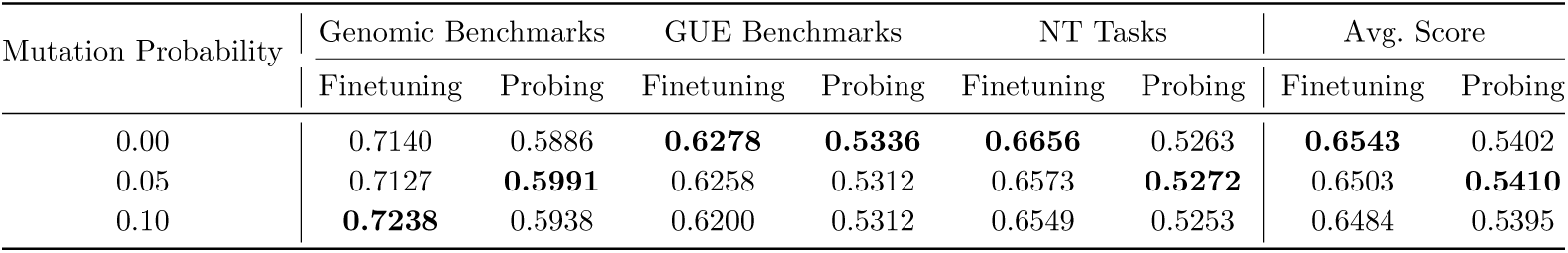
Performance of GenoJEPA with different mutation probabilities across benchmarks. Each task is evaluated through 10-fold cross-validation, and the mean test score is used as the representative result. Average performance across tasks within each benchmark is reported from these mean scores. *Avg Score* denotes the overall average across 55 tasks from three benchmarks. The optimal results are marked in bold.

**Table S11.**
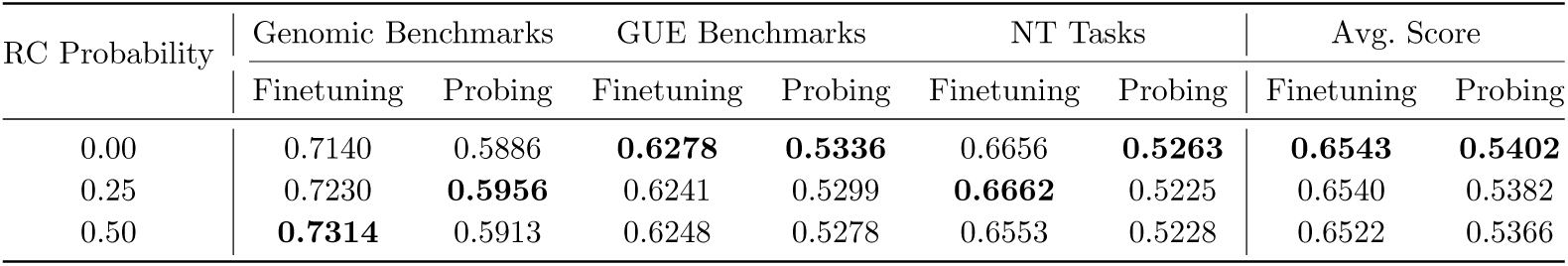
Performance of GenoJEPA with different reverse-complement probabilities across benchmarks. Each task is evaluated through 10-fold cross-validation, and the mean test score is used as the representative result. Average performance across tasks within each benchmark is reported from these mean scores. *Avg Score* denotes the overall average across 55 tasks from three benchmarks. The optimal results are marked in bold.

**Table S12.** Performance of GenoJEPA with different patch sizes across benchmarks. Each task is evaluated through 10-fold cross-validation, and the mean test score is used as the representative result. Average performance across tasks within each benchmark is reported from these mean scores. *Avg Score* denotes the overall average across 55 tasks from three benchmarks. The optimal results are marked in bold.

| Patch Size | Genomic Benchmarks |  | GUE Benchmarks |  | NT Tasks |  | Avg. Score |  |
| --- | --- | --- | --- | --- | --- | --- | --- | --- |
|  | Finetuning | Probing | Finetuning | Probing | Finetuning | Probing | Finetuning | Probing |
| 8 | **0.7340** | **0.6039** | **0.6329** | **0.5354** | 0.6641 | **0.5345** | **0.6597** | **0.5463** |
| 16 | 0.7140 | 0.5886 | 0.6278 | 0.5336 | 0.6656 | 0.5263 | 0.6543 | 0.5402 |
| 32 | 0.7143 | 0.6008 | 0.6227 | 0.5250 | **0.6684** | 0.5238 | 0.6526 | 0.5370 |

**Table S13.** Performance of GenoJEPA with different sliding steps across benchmarks. Each task is evaluated through 10-fold cross-validation, and the mean test score is used as the representative result. Average performance across tasks within each benchmark is reported from these mean scores. *Avg Score* denotes the overall average across 55 tasks from three benchmarks. The optimal results are marked in bold.

| Sliding Step | Genomic Benchmarks |  | GUE Benchmarks |  | NT Tasks |  | Avg. Score |  |
| --- | --- | --- | --- | --- | --- | --- | --- | --- |
|  | Finetuning | Probing | Finetuning | Probing | Finetuning | Probing | Finetuning | Probing |
| 4 | **0.7381** | 0.6034 | 0.6290 | 0.5325 | 0.6558 | **0.5299** | 0.6556 | 0.5433 |
| 8 | 0.7279 | **0.6078** | **0.6327** | 0.5332 | 0.6611 | 0.5296 | **0.6576** | **0.5442** |
| 16 | 0.7140 | 0.5886 | 0.6278 | **0.5336** | **0.6656** | 0.5263 | 0.6543 | 0.5402 |

**Table S14.** Performance of GenoJEPA with different embedding sizes across benchmarks. Each task is evaluated through 10-fold cross-validation, and the mean test score is used as the representative result. Average performance across tasks within each benchmark is reported from these mean scores. *Avg Score* denotes the overall average across 55 tasks from three benchmarks. The optimal results are marked in bold.

| Embedding Size | Genomic Benchmarks |  | GUE Benchmarks |  | NT Tasks |  | Avg. Score |  |
| --- | --- | --- | --- | --- | --- | --- | --- | --- |
|  | Finetuning | Probing | Finetuning | Probing | Finetuning | Probing | Finetuning | Probing |
| 8 | 0.7155 | 0.5954 | 0.6219 | 0.5275 | **0.6706** | 0.5261 | 0.6531 | 0.5382 |
| 16 | 0.7140 | 0.5886 | **0.6278** | 0.5336 | 0.6656 | **0.5263** | **0.6543** | **0.5402** |
| 32 | **0.7191** | **0.5955** | 0.6218 | **0.5361** | 0.6599 | 0.5167 | 0.6502 | 0.5395 |

**Table S15.** Performance of GenoJEPA with different pooling strategies during pretraining across benchmarks. Each task is evaluated through 10-fold cross-validation, and the mean test score is used as the representative result. Average performance across tasks within each benchmark is reported from these mean scores. *Avg Score* denotes the overall average across 55 tasks from three benchmarks. The optimal results are marked in bold.

| Pooling Strategy | Genomic Benchmarks |  | GUE Benchmarks |  | NT Tasks |  | Avg. Score |  |
| --- | --- | --- | --- | --- | --- | --- | --- | --- |
|  | Finetuning | Probing | Finetuning | Probing | Finetuning | Probing | Finetuning | Probing |
| Cls | **0.7263** | 0.5870 | 0.6252 | 0.5315 | 0.6568 | 0.5159 | 0.6521 | 0.5355 |
| Avg | 0.7140 | **0.5886** | **0.6278** | **0.5336** | **0.6656** | **0.5263** | **0.6543** | **0.5402** |
| Max | 0.7130 | 0.5855 | 0.6171 | 0.5138 | 0.6604 | 0.5176 | 0.6470 | 0.5268 |

**Table S16.** Performance of GenoJEPA with different normalization strategies across benchmarks. Each task is evaluated through 10-fold cross-validation, and the mean test score is used as the representative result. Average performance across tasks within each benchmark is reported from these mean scores. *Avg Score* denotes the overall average across 55 tasks from three benchmarks. The optimal results are marked in bold.

| Normalization Strategy | Genomic Benchmarks |  | GUE Benchmarks |  | NT Tasks |  | Avg. Score |  |
| --- | --- | --- | --- | --- | --- | --- | --- | --- |
|  | Finetuning | Probing | Finetuning | Probing | Finetuning | Probing | Finetuning | Probing |
| No Norm | 0.7133 | 0.5865 | 0.6204 | 0.5204 | 0.6616 | 0.5180 | 0.6491 | 0.5304 |
| Layer Norm | 0.7060 | **0.5912** | 0.6224 | 0.5257 | **0.6733** | **0.5332** | 0.6527 | 0.5389 |
| Batch Norm | **0.7140** | 0.5886 | **0.6278** | **0.5336** | 0.6656 | 0.5263 | **0.6543** | **0.5402** |

**Table S17.** Performance of GenoJEPA with different projection strategies across benchmarks. Each task is evaluated through 10-fold cross-validation, and the mean test score is used as the representative result. Average performance across tasks within each benchmark is reported from these mean scores. *Avg Score* denotes the overall average across 55 tasks from three benchmarks. The optimal results are marked in bold.

| Projection Strategy | Genomic Benchmarks |  | GUE Benchmarks |  | NT Tasks |  | Avg. Score |  |
| --- | --- | --- | --- | --- | --- | --- | --- | --- |
|  | Finetuning | Probing | Finetuning | Probing | Finetuning | Probing | Finetuning | Probing |
| No | 0.6871 | 0.5598 | 0.5781 | 0.4822 | 0.6435 | 0.4955 | 0.6174 | 0.4993 |
| Yes | **0.7140** | **0.5886** | **0.6278** | **0.5336** | **0.6656** | **0.5263** | **0.6543** | **0.5402** |

**Table S18.** Performance of GenoJEPA with different projection sizes across benchmarks. Each task is evaluated through 10-fold cross-validation, and the mean test score is used as the representative result. Average performance across tasks within each benchmark is reported from these mean scores. *Avg Score* denotes the overall average across 55 tasks from three benchmarks. The optimal results are marked in bold.

| Projection Size | Genomic Benchmarks |  | GUE Benchmarks |  | NT Tasks |  | Avg. Score |  |
| --- | --- | --- | --- | --- | --- | --- | --- | --- |
|  | Finetuning | Probing | Finetuning | Probing | Finetuning | Probing | Finetuning | Probing |
| 8 | 0.7143 | **0.5948** | 0.6244 | 0.5304 | **0.6686** | 0.5193 | 0.6536 | 0.5373 |
| 16 | 0.7140 | 0.5886 | **0.6278** | **0.5336** | 0.6656 | **0.5263** | **0.6543** | **0.5402** |
| 32 | **0.7159** | 0.5897 | 0.6242 | 0.5254 | 0.6625 | 0.5249 | 0.6517 | 0.5357 |

**Table S19.** Performance of GenoJEPA with different invariance criteria across benchmarks. Each task is evaluated through 10-fold cross-validation, and the mean test score is used as the representative result. Average performance across tasks within each benchmark is reported from these mean scores. *Avg Score* denotes the overall average across 55 tasks from three benchmarks. The optimal results are marked in bold.

| Invariance Criterion | Genomic Benchmarks |  | GUE Benchmarks |  | NT Tasks |  | Avg. Score |  |
| --- | --- | --- | --- | --- | --- | --- | --- | --- |
|  | Finetuning | Probing | Finetuning | Probing | Finetuning | Probing | Finetuning | Probing |
| L1 | 0.6813 | 0.5490 | 0.5770 | 0.4669 | 0.6461 | 0.4740 | 0.6167 | 0.4827 |
| L2 | 0.7140 | 0.5886 | **0.6278** | **0.5336** | 0.6656 | 0.5263 | **0.6543** | **0.5402** |
| Smooth L1 | 0.7184 | **0.5921** | 0.6207 | 0.5270 | **0.6736** | 0.5232 | 0.6540 | 0.5364 |
| Cosine Similarity | **0.7194** | 0.5854 | 0.6235 | 0.5216 | 0.6629 | **0.5265** | 0.6521 | 0.5337 |

**Table S20.**
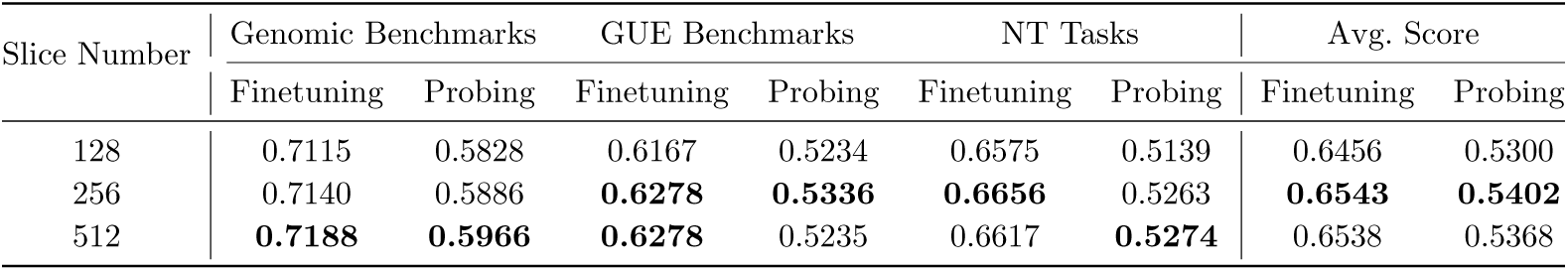
Performance of GenoJEPA with different slice numbers across benchmarks. Each task is evaluated through 10-fold cross-validation, and the mean test score is used as the representative result. Average performance across tasks within each benchmark is reported from these mean scores. *Avg Score* denotes the overall average across 55 tasks from three benchmarks. The optimal results are marked in bold.

**Table S21.** Performance of GenoJEPA with different point numbers across benchmarks. Each task is evaluated through 10-fold cross-validation, and the mean test score is used as the representative result. Average performance across tasks within each benchmark is reported from these mean scores. *Avg Score* denotes the overall average across 55 tasks from three benchmarks. The optimal results are marked in bold.

| Point Number | Genomic Benchmarks |  | GUE Benchmarks |  | NT Tasks |  | Avg. Score |  |
| --- | --- | --- | --- | --- | --- | --- | --- | --- |
|  | Finetuning | Probing | Finetuning | Probing | Finetuning | Probing | Finetuning | Probing |
| 5 | 0.7152 | 0.5823 | 0.6158 | 0.5181 | 0.6445 | 0.5121 | 0.6415 | 0.5266 |
| 17 | 0.7140 | 0.5886 | **0.6278** | **0.5336** | **0.6656** | **0.5263** | **0.6543** | **0.5402** |
| 41 | **0.7173** | **0.5993** | 0.6261 | 0.5287 | 0.6630 | 0.5190 | 0.6531 | 0.5371 |

**Table S22.** Performance of GenoJEPA with different test domains across benchmarks. Each task is evaluated through 10-fold cross-validation, and the mean test score is used as the representative result. Average performance across tasks within each benchmark is reported from these mean scores. *Avg Score* denotes the overall average across 55 tasks from three benchmarks. The optimal results are marked in bold.

| Test Domain | Genomic Benchmarks |  | GUE Benchmarks |  | NT Tasks |  | Avg. Score |  |
| --- | --- | --- | --- | --- | --- | --- | --- | --- |
|  | Finetuning | Probing | Finetuning | Probing | Finetuning | Probing | Finetuning | Probing |
| [-1, 1] | 0.7170 | 0.5806 | 0.6166 | 0.5269 | 0.6534 | 0.5102 | 0.6451 | 0.5302 |
| [-3, 3] | 0.7140 | 0.5886 | 0.6278 | **0.5336** | **0.6656** | 0.5263 | **0.6543** | **0.5402** |
| [-5, 5] | **0.7184** | **0.5955** | **0.6293** | 0.5263 | 0.6605 | **0.5279** | 0.6541 | 0.5381 |

**Table S23.** Performance of GenoJEPA with different SIGReg weights across benchmarks. Each task is evaluated through 10-fold cross-validation, and the mean test score is used as the representative result. Average performance across tasks within each benchmark is reported from these mean scores. *Avg Score* denotes the overall average across 55 tasks from three benchmarks. The optimal results are marked in bold.

| SIGReg Weight | Genomic Benchmarks |  | GUE Benchmarks |  | NT Tasks |  | Avg. Score |  |
| --- | --- | --- | --- | --- | --- | --- | --- | --- |
|  | Finetuning | Probing | Finetuning | Probing | Finetuning | Probing | Finetuning | Probing |
| 0.02 | 0.7129 | 0.5827 | 0.6189 | 0.5206 | 0.6562 | 0.5184 | 0.6465 | 0.5300 |
| 0.05 | 0.7140 | 0.5886 | **0.6278** | **0.5336** | **0.6656** | **0.5263** | **0.6543** | **0.5402** |
| 0.10 | **0.7183** | **0.5899** | 0.6265 | 0.5285 | 0.6569 | 0.5207 | 0.6515 | 0.5360 |

**Table S24.** Performance of GenoJEPA with different learning rates across benchmarks. Each task is evaluated through 10-fold cross-validation, and the mean test score is used as the representative result. Average performance across tasks within each benchmark is reported from these mean scores. *Avg Score* denotes the overall average across 55 tasks from three benchmarks. The optimal results are marked in bold.

| Learning Rate | Genomic Benchmarks |  | GUE Benchmarks |  | NT Tasks |  | Avg. Score |  |
| --- | --- | --- | --- | --- | --- | --- | --- | --- |
|  | Finetuning | Probing | Finetuning | Probing | Finetuning | Probing | Finetuning | Probing |
| $1e^{-3} \rightarrow 1e^{-4}$ | 0.7128 | **0.5894** | 0.6268 | 0.5196 | 0.6577 | 0.5215 | 0.6509 | 0.5317 |
| $5e^{-4} \rightarrow 5e^{-5}$ | 0.7140 | 0.5886 | **0.6278** | **0.5336** | **0.6656** | **0.5263** | **0.6543** | **0.5402** |
| $1e^{-4} \rightarrow 1e^{-5}$ | **0.7191** | 0.5813 | 0.6127 | 0.5246 | 0.6538 | 0.5105 | 0.6436 | 0.5293 |

**Table S25.**
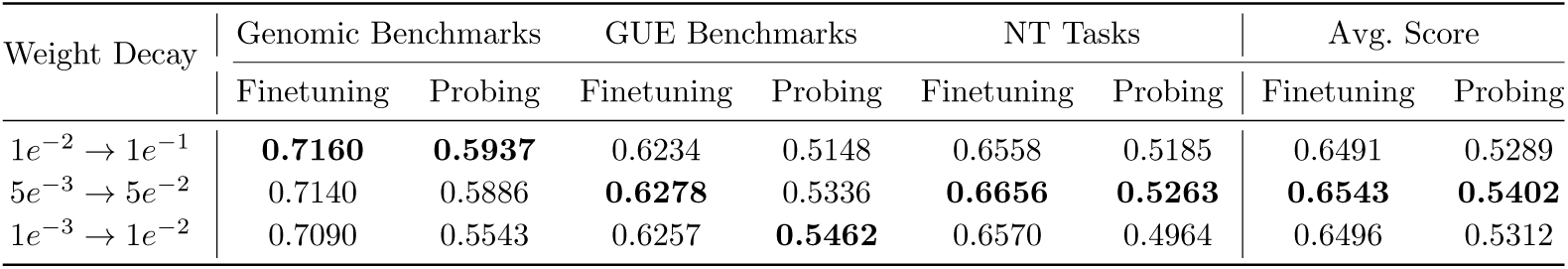
Performance of GenoJEPA with different weight decay values across benchmarks. Each task is evaluated through 10-fold cross-validation, and the mean test score is used as the representative result. Average performance across tasks within each benchmark is reported from these mean scores. *Avg Score* denotes the overall average across 55 tasks from three benchmarks. The optimal results are marked in bold.

| Weight Decay | Genomic Benchmarks |  | GUE Benchmarks |  | NT Tasks |  | Avg. Score |  |
| --- | --- | --- | --- | --- | --- | --- | --- | --- |
|  | Finetuning | Probing | Finetuning | Probing | Finetuning | Probing | Finetuning | Probing |
| $1e^{-2} \rightarrow 1e^{-1}$ | **0.7160** | **0.5937** | 0.6234 | 0.5148 | 0.6558 | 0.5185 | 0.6491 | 0.5289 |
| $5e^{-3} \rightarrow 5e^{-2}$ | 0.7140 | 0.5886 | **0.6278** | 0.5336 | **0.6656** | **0.5263** | **0.6543** | **0.5402** |
| $1e^{-3} \rightarrow 1e^{-2}$ | 0.7090 | 0.5543 | 0.6257 | **0.5462** | 0.6570 | 0.4964 | 0.6496 | 0.5312 |

**Table S26.**
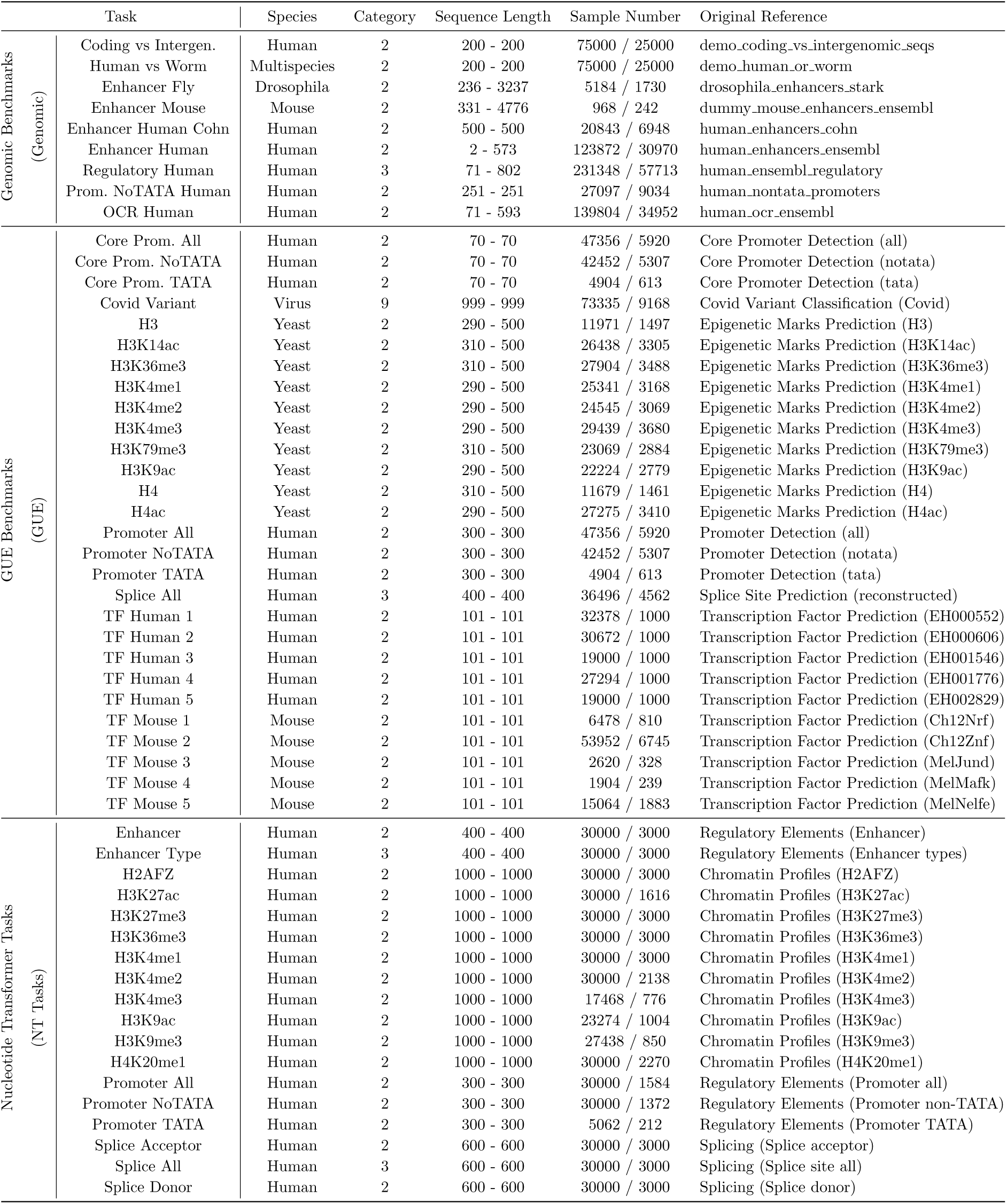
Summary statistics for each task. Some names were shortened in this study for brevity. *Original Reference* denotes the corresponding name in the source paper.

**Table S27.** Finetuning hyperparameters of GenoJEPA and the baselines for each task.

| Task |  | HyenaDNA |  | CaduceusPh |  | DNABERT-2 |  | GROVER |  | NT-v2 |  | GenoJEPA-T |  | GenoJEPA-B |  |
| --- | --- | --- | --- | --- | --- | --- | --- | --- | --- | --- | --- | --- | --- | --- | --- |
|  |  | BS | LR | BS | LR | BS | LR | BS | LR | BS | LR | BS | LR | BS | LR |
| Genomic Benchmarks | Coding vs Intergen. | 64 | $1e^{-4}$ | 128 | $5e^{-5}$ | 64 | $5e^{-5}$ | 128 | $1e^{-3}$ | 64 | $5e^{-5}$ | 256 | $5e^{-4}$ | 256 | $5e^{-5}$ |
| | Human vs Worm | 256 | $1e^{-3}$ | 128 | $5e^{-5}$ | 64 | $1e^{-3}$ | 64 | $1e^{-3}$ | 256 | $5e^{-5}$ | 256 | $1e^{-4}$ | 128 | $5e^{-5}$ |
| | Enhancer Fly | 256 | $5e^{-4}$ | 256 | $5e^{-5}$ | 64 | $1e^{-3}$ | 256 | $1e^{-3}$ | 64 | $1e^{-3}$ | 256 | $5e^{-4}$ | 64 | $1e^{-4}$ |
| | Enhancer Mouse | 256 | $1e^{-3}$ | 128 | $5e^{-5}$ | 64 | $1e^{-4}$ | 64 | $5e^{-4}$ | 256 | $1e^{-4}$ | 64 | $5e^{-4}$ | 64 | $1e^{-4}$ |
| | Enhancer Human Cohn | 256 | $5e^{-5}$ | 128 | $1e^{-3}$ | 128 | $5e^{-5}$ | 64 | $5e^{-5}$ | 64 | $5e^{-5}$ | 256 | $5e^{-5}$ | 128 | $1e^{-3}$ |
| | Enhancer Human | 256 | $5e^{-4}$ | 64 | $1e^{-4}$ | 64 | $5e^{-5}$ | 256 | $5e^{-4}$ | 64 | $5e^{-5}$ | 256 | $1e^{-3}$ | 256 | $1e^{-4}$ |
| | Regulatory Human | 256 | $1e^{-3}$ | 128 | $1e^{-4}$ | 64 | $1e^{-3}$ | 256 | $5e^{-4}$ | 256 | $1e^{-3}$ | 256 | $1e^{-4}$ | 64 | $5e^{-4}$ |
| | Prom. NoTATA Human | 256 | $5e^{-4}$ | 128 | $5e^{-5}$ | 64 | $1e^{-3}$ | 64 | $5e^{-4}$ | 256 | $5e^{-5}$ | 256 | $5e^{-4}$ | 128 | $1e^{-4}$ |
| | OCR Human | 256 | $5e^{-5}$ | 256 | $5e^{-5}$ | 128 | $1e^{-4}$ | 64 | $1e^{-3}$ | 64 | $5e^{-4}$ | 64 | $1e^{-3}$ | 256 | $5e^{-4}$ |
| GUE Benchmarks | Core Prom. All | 256 | $5e^{-4}$ | 64 | $1e^{-3}$ | 64 | $5e^{-5}$ | 128 | $5e^{-5}$ | 256 | $5e^{-5}$ | 64 | $5e^{-4}$ | 128 | $1e^{-3}$ |
| | Core Prom. NoTATA | 256 | $1e^{-3}$ | 128 | $5e^{-5}$ | 64 | $1e^{-4}$ | 128 | $1e^{-3}$ | 256 | $1e^{-3}$ | 64 | $5e^{-4}$ | 256 | $5e^{-4}$ |
| | Core Prom. TATA | 256 | $5e^{-4}$ | 64 | $1e^{-3}$ | 64 | $1e^{-4}$ | 256 | $1e^{-3}$ | 256 | $5e^{-5}$ | 128 | $1e^{-3}$ | 64 | $5e^{-4}$ |
| | Covid Variant | 128 | $1e^{-3}$ | 256 | $5e^{-5}$ | 64 | $1e^{-4}$ | 64 | $1e^{-3}$ | 64 | $5e^{-5}$ | 256 | $5e^{-4}$ | 64 | $5e^{-5}$ |
| | H3 | 256 | $5e^{-5}$ | 128 | $1e^{-4}$ | 128 | $1e^{-3}$ | 128 | $5e^{-5}$ | 64 | $1e^{-3}$ | 256 | $1e^{-3}$ | 128 | $1e^{-4}$ |
| | H3K14ac | 256 | $1e^{-4}$ | 256 | $1e^{-4}$ | 64 | $1e^{-3}$ | 64 | $5e^{-4}$ | 256 | $5e^{-5}$ | 64 | $5e^{-4}$ | 64 | $1e^{-4}$ |
| | H3K36me3 | 256 | $1e^{-3}$ | 128 | $1e^{-4}$ | 64 | $5e^{-5}$ | 128 | $1e^{-3}$ | 256 | $5e^{-4}$ | 64 | $1e^{-3}$ | 128 | $1e^{-3}$ |
| | H3K4me1 | 256 | $5e^{-4}$ | 64 | $5e^{-5}$ | 64 | $1e^{-4}$ | 64 | $5e^{-4}$ | 64 | $5e^{-5}$ | 64 | $1e^{-3}$ | 128 | $5e^{-4}$ |
| | H3K4me2 | 256 | $5e^{-4}$ | 128 | $5e^{-5}$ | 64 | $5e^{-5}$ | 256 | $5e^{-4}$ | 128 | $1e^{-4}$ | 64 | $5e^{-4}$ | 256 | $1e^{-4}$ |
| | H3K4me3 | 256 | $5e^{-4}$ | 256 | $1e^{-4}$ | 64 | $1e^{-4}$ | 128 | $1e^{-3}$ | 256 | $5e^{-5}$ | 64 | $1e^{-3}$ | 256 | $5e^{-4}$ |
| | H3K79me3 | 256 | $1e^{-3}$ | 128 | $5e^{-5}$ | 64 | $1e^{-3}$ | 64 | $1e^{-3}$ | 64 | $1e^{-3}$ | 64 | $5e^{-4}$ | 256 | $1e^{-4}$ |
| | H3K9ac | 256 | $5e^{-4}$ | 256 | $5e^{-5}$ | 128 | $1e^{-4}$ | 64 | $1e^{-4}$ | 64 | $1e^{-4}$ | 64 | $1e^{-4}$ | 64 | $1e^{-4}$ |
| | H4 | 256 | $1e^{-3}$ | 128 | $1e^{-3}$ | 128 | $1e^{-3}$ | 64 | $5e^{-4}$ | 256 | $5e^{-5}$ | 128 | $5e^{-5}$ | 256 | $5e^{-4}$ |
| | H4ac | 256 | $1e^{-3}$ | 256 | $1e^{-4}$ | 128 | $1e^{-3}$ | 64 | $1e^{-3}$ | 256 | $5e^{-5}$ | 256 | $1e^{-3}$ | 64 | $1e^{-3}$ |
| | Promoter All | 256 | $1e^{-4}$ | 256 | $1e^{-4}$ | 64 | $1e^{-3}$ | 64 | $1e^{-3}$ | 64 | $1e^{-4}$ | 256 | $5e^{-4}$ | 64 | $5e^{-4}$ |
| | Promoter NoTATA | 256 | $5e^{-5}$ | 128 | $5e^{-5}$ | 64 | $5e^{-5}$ | 64 | $5e^{-5}$ | 64 | $1e^{-3}$ | 128 | $5e^{-5}$ | 128 | $1e^{-4}$ |
| | Promoter TATA | 256 | $5e^{-4}$ | 256 | $1e^{-3}$ | 64 | $5e^{-4}$ | 64 | $5e^{-4}$ | 256 | $1e^{-3}$ | 64 | $5e^{-4}$ | 256 | $1e^{-3}$ |
| | Splice All | 256 | $5e^{-5}$ | 64 | $5e^{-5}$ | 128 | $1e^{-4}$ | 256 | $1e^{-3}$ | 256 | $1e^{-4}$ | 256 | $5e^{-5}$ | 64 | $5e^{-5}$ |
| | TF Human 1 | 256 | $1e^{-3}$ | 256 | $5e^{-5}$ | 64 | $1e^{-3}$ | 128 | $1e^{-3}$ | 256 | $5e^{-5}$ | 256 | $5e^{-5}$ | 64 | $1e^{-4}$ |
| | TF Human 2 | 256 | $5e^{-5}$ | 64 | $1e^{-3}$ | 128 | $1e^{-3}$ | 256 | $5e^{-5}$ | 256 | $1e^{-4}$ | 128 | $5e^{-5}$ | 256 | $1e^{-4}$ |
| | TF Human 3 | 64 | $1e^{-3}$ | 256 | $5e^{-5}$ | 128 | $1e^{-3}$ | 128 | $5e^{-5}$ | 256 | $1e^{-3}$ | 256 | $1e^{-3}$ | 64 | $5e^{-4}$ |
| | TF Human 4 | 256 | $5e^{-5}$ | 256 | $1e^{-4}$ | 64 | $1e^{-3}$ | 128 | $1e^{-3}$ | 256 | $5e^{-5}$ | 128 | $1e^{-3}$ | 128 | $1e^{-3}$ |
| | TF Human 5 | 256 | $1e^{-3}$ | 128 | $5e^{-5}$ | 64 | $1e^{-4}$ | 64 | $1e^{-3}$ | 64 | $1e^{-3}$ | 256 | $1e^{-3}$ | 128 | $1e^{-3}$ |
| | TF Mouse 1 | 256 | $5e^{-4}$ | 64 | $5e^{-5}$ | 128 | $1e^{-4}$ | 64 | $5e^{-5}$ | 256 | $5e^{-5}$ | 64 | $5e^{-4}$ | 64 | $5e^{-4}$ |
| | TF Mouse 2 | 256 | $1e^{-3}$ | 64 | $1e^{-4}$ | 64 | $5e^{-5}$ | 128 | $1e^{-3}$ | 128 | $5e^{-5}$ | 256 | $1e^{-3}$ | 64 | $1e^{-4}$ |
| | TF Mouse 3 | 256 | $5e^{-4}$ | 256 | $5e^{-5}$ | 128 | $5e^{-4}$ | 64 | $5e^{-5}$ | 256 | $5e^{-5}$ | 256 | $1e^{-3}$ | 128 | $1e^{-3}$ |
| | TF Mouse 4 | 256 | $5e^{-4}$ | 64 | $5e^{-5}$ | 64 | $5e^{-5}$ | 64 | $1e^{-4}$ | 64 | $5e^{-5}$ | 256 | $5e^{-4}$ | 128 | $5e^{-4}$ |
| | TF Mouse 5 | 128 | $5e^{-4}$ | 128 | $5e^{-5}$ | 128 | $5e^{-5}$ | 256 | $1e^{-4}$ | 256 | $1e^{-4}$ | 128 | $5e^{-4}$ | 256 | $1e^{-4}$ |
| Nucleotide Transformer Tasks | Enhancer | 256 | $1e^{-3}$ | 64 | $5e^{-5}$ | 256 | $1e^{-3}$ | 256 | $1e^{-3}$ | 128 | $1e^{-3}$ | 64 | $5e^{-4}$ | 64 | $1e^{-3}$ |
| | Enhancer Type | 256 | $1e^{-3}$ | 64 | $5e^{-5}$ | 64 | $1e^{-4}$ | 256 | $1e^{-3}$ | 128 | $5e^{-4}$ | 64 | $1e^{-3}$ | 128 | $1e^{-3}$ |
| | H2AFZ | 256 | $5e^{-4}$ | 256 | $5e^{-5}$ | 64 | $1e^{-3}$ | 128 | $1e^{-3}$ | 256 | $5e^{-5}$ | 128 | $5e^{-4}$ | 256 | $1e^{-3}$ |
| | H3K27ac | 256 | $1e^{-3}$ | 256 | $1e^{-4}$ | 64 | $1e^{-4}$ | 64 | $5e^{-4}$ | 64 | $1e^{-4}$ | 256 | $1e^{-4}$ | 256 | $5e^{-4}$ |
| | H3K27me3 | 256 | $5e^{-4}$ | 256 | $1e^{-4}$ | 64 | $5e^{-5}$ | 64 | $5e^{-5}$ | 128 | $1e^{-4}$ | 64 | $1e^{-3}$ | 256 | $1e^{-4}$ |
| | H3K36me3 | 256 | $5e^{-5}$ | 128 | $5e^{-5}$ | 64 | $1e^{-4}$ | 64 | $5e^{-5}$ | 128 | $5e^{-5}$ | 256 | $5e^{-5}$ | 64 | $1e^{-4}$ |
| | H3K4me1 | 256 | $1e^{-3}$ | 128 | $5e^{-5}$ | 64 | $5e^{-5}$ | 64 | $1e^{-3}$ | 64 | $1e^{-4}$ | 256 | $5e^{-4}$ | 64 | $1e^{-4}$ |
| | H3K4me2 | 256 | $5e^{-4}$ | 256 | $5e^{-5}$ | 64 | $1e^{-4}$ | 256 | $1e^{-3}$ | 128 | $5e^{-5}$ | 128 | $5e^{-4}$ | 64 | $5e^{-5}$ |
| | H3K4me3 | 256 | $1e^{-4}$ | 64 | $1e^{-3}$ | 128 | $5e^{-4}$ | 128 | $1e^{-4}$ | 64 | $5e^{-5}$ | 128 | $1e^{-3}$ | 64 | $1e^{-4}$ |
| | H3K9ac | 256 | $5e^{-4}$ | 128 | $5e^{-5}$ | 64 | $5e^{-5}$ | 256 | $5e^{-4}$ | 256 | $5e^{-5}$ | 128 | $5e^{-4}$ | 128 | $5e^{-5}$ |
| | H3K9me3 | 64 | $5e^{-5}$ | 128 | $1e^{-4}$ | 64 | $1e^{-4}$ | 64 | $5e^{-5}$ | 64 | $5e^{-5}$ | 256 | $1e^{-3}$ | 64 | $5e^{-4}$ |
| | H4K20me1 | 256 | $5e^{-4}$ | 256 | $5e^{-5}$ | 128 | $1e^{-3}$ | 64 | $5e^{-5}$ | 64 | $1e^{-4}$ | 256 | $1e^{-4}$ | 128 | $5e^{-5}$ |
| | Promoter All | 256 | $1e^{-3}$ | 128 | $5e^{-5}$ | 128 | $1e^{-3}$ | 64 | $5e^{-5}$ | 64 | $1e^{-4}$ | 128 | $5e^{-4}$ | 256 | $5e^{-4}$ |
| | Promoter NoTATA | 64 | $5e^{-4}$ | 64 | $5e^{-5}$ | 64 | $5e^{-5}$ | 128 | $1e^{-3}$ | 64 | $1e^{-3}$ | 64 | $1e^{-3}$ | 256 | $1e^{-3}$ |
| | Promoter TATA | 256 | $1e^{-3}$ | 64 | $1e^{-4}$ | 64 | $1e^{-4}$ | 128 | $1e^{-4}$ | 256 | $5e^{-5}$ | 128 | $1e^{-3}$ | 64 | $1e^{-4}$ |
| | Splice Acceptor | 128 | $5e^{-5}$ | 256 | $1e^{-4}$ | 64 | $1e^{-3}$ | 128 | $1e^{-3}$ | 64 | $5e^{-5}$ | 128 | $5e^{-5}$ | 64 | $5e^{-4}$ |
| | Splice All | 64 | $5e^{-4}$ | 64 | $1e^{-4}$ | 128 | $5e^{-5}$ | 64 | $5e^{-5}$ | 256 | $5e^{-5}$ | 64 | $1e^{-3}$ | 256 | $1e^{-3}$ |
| | Splice Donor | 256 | $5e^{-4}$ | 64 | $1e^{-4}$ | 64 | $1e^{-3}$ | 64 | $5e^{-5}$ | 256 | $5e^{-5}$ | 256 | $5e^{-4}$ | 256 | $5e^{-4}$ |

**Table S28.** Probing performance of GenoJEPA and the baselines for each task. Each task is evaluated through 10-fold cross-validation, and the mean and standard deviation of the test scores across folds are reported as the representative results. *Avg Score* and *Avg Rank* denote the averages across 55 tasks from three benchmarks. The optimal and second optimal values are marked in bold and underlined, respectively.

| Task |  | HyenaDNA<br>(7M) | CaduceusPh<br>(8M) | DNABERT-2<br>(117M) | GROVER<br>(87M) | NT-v2<br>(494M) | GenoJEPA-T<br>(6M) | GenoJEPA-B<br>(52M) |
| --- | --- | --- | --- | --- | --- | --- | --- | --- |
| Genomic Benchmarks | Coding vs Intergen. | 0.695 ± 0.020 | 0.738 ± 0.022 | 0.819 ± 0.010 | 0.818 ± 0.012 | 0.768 ± 0.010 | <u>0.847 ± 0.009</u> | **0.886 ± 0.005** |
|  | Human vs Worm | 0.793 ± 0.016 | 0.689 ± 0.015 | 0.904 ± 0.017 | 0.892 ± 0.016 | 0.891 ± 0.006 | <u>0.927 ± 0.016</u> | **0.946 ± 0.003** |
|  | Enhancer Fly | 0.379 ± 0.003 | 0.120 ± 0.016 | <u>0.444 ± 0.009</u> | 0.385 ± 0.011 | 0.436 ± 0.004 | 0.419 ± 0.002 | **0.489 ± 0.003** |
|  | Enhancer Mouse | 0.441 ± 0.018 | 0.351 ± 0.007 | 0.634 ± 0.017 | 0.566 ± 0.012 | **0.733 ± 0.009** | 0.632 ± 0.016 | <u>0.677 ± 0.004</u> |
|  | Enhancer Human Cohn | 0.438 ± 0.004 | 0.420 ± 0.006 | <u>0.506 ± 0.014</u> | 0.466 ± 0.019 | 0.445 ± 0.012 | 0.488 ± 0.005 | **0.531 ± 0.002** |
|  | Enhancer Human | 0.346 ± 0.011 | 0.382 ± 0.005 | 0.504 ± 0.006 | 0.490 ± 0.009 | <u>0.528 ± 0.009</u> | 0.490 ± 0.008 | **0.531 ± 0.005** |
|  | Regulatory Human | **0.966 ± 0.002** | 0.428 ± 0.003 | 0.507 ± 0.009 | 0.458 ± 0.007 | 0.490 ± 0.005 | 0.446 ± 0.008 | <u>0.534 ± 0.003</u> |
|  | Prom. NoTATA Human | 0.649 ± 0.022 | 0.530 ± 0.016 | 0.684 ± 0.017 | 0.679 ± 0.014 | 0.673 ± 0.012 | <u>0.690 ± 0.009</u> | **0.725 ± 0.015** |
|  | OCR Human | 0.297 ± 0.021 | 0.229 ± 0.014 | 0.348 ± 0.015 | <u>0.367 ± 0.022</u> | 0.345 ± 0.002 | 0.358 ± 0.004 | **0.413 ± 0.023** |
| GUE Benchmarks | Core Prom. All | 0.519 ± 0.010 | 0.467 ± 0.017 | 0.578 ± 0.011 | 0.567 ± 0.008 | 0.584 ± 0.010 | <u>0.589 ± 0.006</u> | **0.607 ± 0.006** |
|  | Core Prom. NoTATA | 0.570 ± 0.011 | 0.502 ± 0.014 | 0.596 ± 0.020 | 0.612 ± 0.015 | 0.619 ± 0.010 | <u>0.649 ± 0.003</u> | **0.675 ± 0.016** |
|  | Core Prom. TATA | 0.434 ± 0.002 | 0.404 ± 0.004 | 0.516 ± 0.014 | 0.452 ± 0.018 | **0.631 ± 0.008** | <u>0.518 ± 0.002</u> | 0.455 ± 0.003 |
|  | Covid Variant | 0.041 ± 0.011 | 0.031 ± 0.004 | **0.204 ± 0.008** | <u>0.196 ± 0.004</u> | 0.093 ± 0.006 | 0.135 ± 0.005 | 0.136 ± 0.003 |
|  | H3 | 0.660 ± 0.001 | 0.538 ± 0.007 | 0.660 ± 0.015 | 0.660 ± 0.004 | 0.632 ± 0.017 | <u>0.703 ± 0.010</u> | **0.717 ± 0.006** |
|  | H3K14ac | 0.326 ± 0.004 | 0.224 ± 0.021 | 0.401 ± 0.004 | 0.357 ± 0.008 | 0.372 ± 0.007 | <u>0.412 ± 0.008</u> | **0.484 ± 0.010** |
|  | H3K36me3 | 0.394 ± 0.009 | 0.278 ± 0.009 | 0.445 ± 0.004 | 0.387 ± 0.012 | 0.444 ± 0.002 | <u>0.465 ± 0.017</u> | **0.517 ± 0.015** |
|  | H3K4me1 | 0.330 ± 0.011 | 0.265 ± 0.006 | 0.341 ± 0.004 | 0.352 ± 0.011 | 0.352 ± 0.007 | <u>0.386 ± 0.021</u> | **0.432 ± 0.009** |
|  | H3K4me2 | 0.328 ± 0.016 | 0.308 ± 0.005 | 0.304 ± 0.008 | 0.308 ± 0.026 | 0.313 ± 0.006 | <u>0.332 ± 0.006</u> | **0.345 ± 0.006** |
|  | H3K4me3 | 0.202 ± 0.003 | 0.149 ± 0.019 | 0.255 ± 0.010 | 0.245 ± 0.015 | 0.240 ± 0.012 | <u>0.278 ± 0.003</u> | **0.326 ± 0.009** |
|  | H3K79me3 | 0.519 ± 0.008 | 0.364 ± 0.033 | <u>0.590 ± 0.014</u> | 0.521 ± 0.010 | 0.570 ± 0.005 | 0.576 ± 0.004 | **0.612 ± 0.019** |
|  | H3K9ac | 0.433 ± 0.025 | 0.337 ± 0.019 | <u>0.498 ± 0.008</u> | 0.444 ± 0.011 | 0.434 ± 0.005 | 0.494 ± 0.006 | **0.538 ± 0.006** |
|  | H4 | 0.705 ± 0.016 | 0.472 ± 0.004 | 0.745 ± 0.013 | 0.728 ± 0.016 | 0.745 ± 0.011 | **0.797 ± 0.004** | <u>0.789 ± 0.010</u> |
|  | H4ac | 0.314 ± 0.013 | 0.217 ± 0.016 | 0.358 ± 0.005 | 0.312 ± 0.008 | 0.318 ± 0.006 | <u>0.370 ± 0.002</u> | **0.420 ± 0.008** |
|  | Promoter All | 0.743 ± 0.019 | 0.577 ± 0.013 | 0.788 ± 0.021 | 0.769 ± 0.011 | 0.777 ± 0.007 | <u>0.795 ± 0.013</u> | **0.826 ± 0.005** |
|  | Promoter NoTATA | 0.825 ± 0.009 | 0.638 ± 0.023 | 0.881 ± 0.003 | <u>0.893 ± 0.008</u> | 0.879 ± 0.017 | 0.888 ± 0.013 | **0.927 ± 0.007** |
|  | Promoter TATA | <u>0.568 ± 0.034</u> | 0.280 ± 0.004 | 0.412 ± 0.005 | 0.476 ± 0.011 | 0.527 ± 0.013 | 0.453 ± 0.010 | **0.620 ± 0.002** |
|  | Splice All | 0.337 ± 0.013 | 0.327 ± 0.011 | 0.353 ± 0.009 | 0.350 ± 0.012 | 0.365 ± 0.015 | <u>0.408 ± 0.011</u> | **0.511 ± 0.003** |
|  | TF Human 1 | 0.567 ± 0.019 | 0.491 ± 0.015 | 0.606 ± 0.010 | 0.652 ± 0.005 | 0.560 ± 0.011 | <u>0.655 ± 0.009</u> | **0.675 ± 0.003** |
|  | TF Human 2 | 0.622 ± 0.009 | 0.522 ± 0.008 | 0.682 ± 0.012 | 0.684 ± 0.010 | 0.655 ± 0.009 | <u>0.692 ± 0.005</u> | **0.704 ± 0.012** |
|  | TF Human 3 | 0.434 ± 0.018 | 0.380 ± 0.011 | <u>0.550 ± 0.004</u> | 0.516 ± 0.013 | 0.506 ± 0.002 | 0.537 ± 0.005 | **0.560 ± 0.002** |
|  | TF Human 4 | 0.293 ± 0.008 | 0.253 ± 0.014 | 0.352 ± 0.004 | 0.393 ± 0.017 | 0.330 ± 0.011 | <u>0.446 ± 0.007</u> | **0.481 ± 0.011** |
|  | TF Human 5 | 0.582 ± 0.003 | 0.503 ± 0.003 | <u>0.640 ± 0.017</u> | 0.598 ± 0.019 | 0.615 ± 0.003 | 0.623 ± 0.002 | **0.681 ± 0.008** |
|  | TF Mouse 1 | 0.223 ± 0.003 | 0.075 ± 0.017 | 0.389 ± 0.004 | 0.335 ± 0.002 | <u>0.403 ± 0.011</u> | 0.399 ± 0.002 | **0.556 ± 0.007** |
|  | TF Mouse 2 | 0.595 ± 0.021 | 0.502 ± 0.004 | 0.700 ± 0.012 | 0.687 ± 0.014 | 0.712 ± 0.011 | <u>0.760 ± 0.015</u> | **0.838 ± 0.016** |
|  | TF Mouse 3 | 0.492 ± 0.014 | 0.401 ± 0.008 | 0.689 ± 0.015 | 0.667 ± 0.026 | <u>0.711 ± 0.009</u> | 0.694 ± 0.006 | **0.746 ± 0.006** |
|  | TF Mouse 4 | 0.261 ± 0.009 | 0.170 ± 0.020 | 0.406 ± 0.026 | 0.468 ± 0.012 | 0.371 ± 0.011 | <u>0.469 ± 0.004</u> | **0.653 ± 0.012** |
|  | TF Mouse 5 | 0.236 ± 0.003 | 0.072 ± 0.004 | 0.364 ± 0.011 | 0.309 ± 0.012 | 0.323 ± 0.022 | <u>0.415 ± 0.015</u> | **0.513 ± 0.005** |
| Nucleotide Transformer Tasks | Enhancer | 0.459 ± 0.019 | 0.425 ± 0.007 | 0.473 ± 0.009 | <u>0.488 ± 0.009</u> | 0.481 ± 0.004 | 0.482 ± 0.010 | **0.489 ± 0.004** |
|  | Enhancer Type | 0.498 ± 0.010 | 0.480 ± 0.006 | <u>0.519 ± 0.006</u> | 0.510 ± 0.010 | 0.517 ± 0.004 | 0.514 ± 0.017 | **0.532 ± 0.007** |
|  | H2AFZ | **0.475 ± 0.021** | 0.337 ± 0.023 | 0.395 ± 0.009 | 0.403 ± 0.011 | 0.375 ± 0.020 | 0.394 ± 0.007 | <u>0.421 ± 0.008</u> |
|  | H3K27ac | 0.424 ± 0.010 | 0.277 ± 0.008 | 0.431 ± 0.015 | 0.424 ± 0.010 | 0.377 ± 0.012 | <u>0.438 ± 0.008</u> | **0.459 ± 0.002** |
|  | H3K27me3 | 0.539 ± 0.018 | 0.474 ± 0.016 | 0.532 ± 0.019 | 0.537 ± 0.007 | 0.513 ± 0.005 | <u>0.541 ± 0.002</u> | **0.551 ± 0.010** |
|  | H3K36me3 | 0.518 ± 0.010 | 0.355 ± 0.010 | 0.531 ± 0.026 | 0.483 ± 0.005 | 0.494 ± 0.009 | <u>0.533 ± 0.005</u> | **0.623 ± 0.010** |
|  | H3K4me1 | 0.408 ± 0.003 | 0.345 ± 0.016 | <u>0.419 ± 0.004</u> | 0.394 ± 0.006 | 0.402 ± 0.003 | 0.418 ± 0.008 | **0.458 ± 0.002** |
|  | H3K4me2 | **0.502 ± 0.017** | 0.411 ± 0.022 | 0.469 ± 0.012 | 0.464 ± 0.002 | 0.469 ± 0.009 | 0.445 ± 0.011 | <u>0.498 ± 0.016</u> |
|  | H3K4me3 | <u>0.628 ± 0.006</u> | 0.482 ± 0.013 | 0.596 ± 0.004 | 0.575 ± 0.011 | **0.645 ± 0.013** | 0.619 ± 0.003 | 0.613 ± 0.002 |
|  | H3K9ac | 0.474 ± 0.004 | 0.380 ± 0.021 | 0.509 ± 0.016 | 0.477 ± 0.004 | 0.499 ± 0.008 | <u>0.543 ± 0.003</u> | **0.565 ± 0.017** |
|  | H3K9me3 | <u>0.361 ± 0.017</u> | 0.220 ± 0.009 | 0.326 ± 0.013 | 0.360 ± 0.018 | 0.320 ± 0.007 | 0.344 ± 0.014 | **0.363 ± 0.012** |
|  | H4K20me1 | 0.579 ± 0.008 | 0.506 ± 0.018 | 0.578 ± 0.011 | 0.532 ± 0.018 | 0.550 ± 0.001 | <u>0.593 ± 0.015</u> | **0.625 ± 0.003** |
|  | Promoter All | <u>0.716 ± 0.013</u> | 0.684 ± 0.024 | 0.677 ± 0.011 | 0.696 ± 0.003 | 0.715 ± 0.007 | 0.711 ± 0.009 | **0.753 ± 0.004** |
|  | Promoter NoTATA | 0.728 ± 0.011 | 0.718 ± 0.008 | <u>0.744 ± 0.015</u> | 0.725 ± 0.011 | 0.743 ± 0.003 | 0.742 ± 0.013 | **0.790 ± 0.007** |
|  | Promoter TATA | **0.768 ± 0.005** | 0.664 ± 0.011 | 0.687 ± 0.006 | <u>0.762 ± 0.008</u> | 0.710 ± 0.008 | 0.697 ± 0.004 | 0.741 ± 0.008 |
|  | Splice Acceptor | 0.456 ± 0.002 | 0.438 ± 0.019 | <u>0.612 ± 0.018</u> | 0.463 ± 0.018 | 0.467 ± 0.005 | 0.566 ± 0.006 | **0.680 ± 0.006** |
|  | Splice All | 0.270 ± 0.013 | 0.273 ± 0.008 | 0.322 ± 0.017 | 0.281 ± 0.009 | <u>0.330 ± 0.014</u> | 0.322 ± 0.006 | **0.434 ± 0.007** |
|  | Splice Donor | 0.460 ± 0.012 | 0.443 ± 0.016 | <u>0.625 ± 0.004</u> | 0.496 ± 0.003 | 0.521 ± 0.021 | 0.572 ± 0.002 | **0.698 ± 0.008** |
| Avg. Score |  | 0.488 | 0.392 | 0.529 | 0.511 | 0.519 | <u>0.540</u> | **0.589** |
| Avg. Rank |  | 5.036 | 6.909 | 3.491 | 4.400 | 4.018 | <u>2.873</u> | **1.273** |

**Table S29.** Finetuning performance of GenoJEPA and the baselines for each task. Each task is evaluated through 10-fold cross-validation, and the mean and standard deviation of the test scores across folds are reported as the representative results. *Avg Score* and *Avg Rank* denote the averages across 55 tasks from three benchmarks. The optimal and second optimal values are marked in bold and underlined, respectively.

| Task |  | HyenaDNA<br>(7M) | CaduceusPh<br>(8M) | DNABERT-2<br>(117M) | GROVER<br>(87M) | NT-v2<br>(494M) | GenoJEPA-T<br>(6M) | GenoJEPA-B<br>(52M) |
| --- | --- | --- | --- | --- | --- | --- | --- | --- |
| Genomic Benchmarks | Coding vs Intergen. | 0.795 ± 0.022 | 0.851 ± 0.025 | 0.872 ± 0.015 | 0.815 ± 0.029 | <u>0.888 ± 0.008</u> | 0.860 ± 0.011 | **0.908 ± 0.007** |
|  | Human vs Worm | 0.913 ± 0.017 | 0.934 ± 0.023 | <u>0.950 ± 0.015</u> | 0.894 ± 0.019 | <u>0.950 ± 0.007</u> | 0.942 ± 0.017 | **0.963 ± 0.003** |
|  | Enhancer Fly | 0.432 ± 0.004 | 0.513 ± 0.019 | 0.534 ± 0.017 | 0.452 ± 0.011 | **0.615 ± 0.004** | 0.533 ± 0.003 | <u>0.596 ± 0.004</u> |
|  | Enhancer Mouse | 0.513 ± 0.017 | 0.555 ± 0.018 | 0.724 ± 0.016 | 0.507 ± 0.015 | <u>0.734 ± 0.016</u> | 0.662 ± 0.017 | **0.757 ± 0.005** |
|  | Enhancer Human Cohn | 0.408 ± 0.007 | 0.478 ± 0.007 | 0.487 ± 0.024 | 0.482 ± 0.016 | <u>0.516 ± 0.012</u> | 0.489 ± 0.006 | **0.530 ± 0.002** |
|  | Enhancer Human | 0.755 ± 0.016 | 0.775 ± 0.005 | 0.755 ± 0.007 | **0.803 ± 0.010** | <u>0.801 ± 0.015</u> | 0.759 ± 0.011 | 0.788 ± 0.006 |
|  | Regulatory Human | 0.853 ± 0.002 | 0.891 ± 0.004 | 0.828 ± 0.011 | 0.852 ± 0.011 | 0.883 ± 0.008 | <u>0.907 ± 0.008</u> | **0.915 ± 0.003** |
|  | Prom. NoTATA Human | 0.802 ± 0.029 | 0.813 ± 0.022 | **0.904 ± 0.017** | 0.834 ± 0.026 | 0.832 ± 0.015 | 0.779 ± 0.008 | <u>0.870 ± 0.015</u> |
|  | OCR Human | 0.569 ± 0.021 | 0.588 ± 0.019 | <u>0.591 ± 0.015</u> | 0.579 ± 0.018 | **0.594 ± 0.002** | 0.494 ± 0.006 | 0.550 ± 0.022 |
| GUE Benchmarks | Core Prom. All | 0.641 ± 0.014 | 0.645 ± 0.015 | **0.717 ± 0.016** | 0.635 ± 0.014 | 0.694 ± 0.019 | <u>0.705 ± 0.008</u> | 0.703 ± 0.008 |
|  | Core Prom. NoTATA | 0.662 ± 0.017 | 0.664 ± 0.024 | <u>0.712 ± 0.028</u> | 0.653 ± 0.019 | 0.707 ± 0.029 | 0.704 ± 0.006 | **0.725 ± 0.015** |
|  | Core Prom. TATA | 0.709 ± 0.003 | 0.741 ± 0.007 | 0.733 ± 0.016 | 0.584 ± 0.034 | 0.741 ± 0.010 | <u>0.763 ± 0.002</u> | **0.819 ± 0.004** |
|  | Covid Variant | 0.619 ± 0.015 | 0.649 ± 0.006 | 0.650 ± 0.008 | 0.651 ± 0.006 | <u>0.681 ± 0.008</u> | 0.657 ± 0.007 | **0.685 ± 0.006** |
|  | H3 | 0.713 ± 0.002 | 0.734 ± 0.008 | 0.777 ± 0.019 | 0.768 ± 0.005 | **0.790 ± 0.015** | 0.762 ± 0.011 | <u>0.786 ± 0.005</u> |
|  | H3K14ac | 0.404 ± 0.005 | 0.500 ± 0.039 | **0.610 ± 0.004** | 0.423 ± 0.009 | 0.542 ± 0.009 | 0.459 ± 0.014 | <u>0.566 ± 0.011</u> |
|  | H3K36me3 | 0.482 ± 0.013 | 0.573 ± 0.010 | <u>0.624 ± 0.004</u> | 0.503 ± 0.016 | **0.658 ± 0.002** | 0.541 ± 0.021 | 0.621 ± 0.011 |
|  | H3K4me1 | 0.372 ± 0.016 | 0.455 ± 0.013 | 0.476 ± 0.005 | 0.440 ± 0.019 | **0.583 ± 0.011** | 0.450 ± 0.021 | <u>0.542 ± 0.011</u> |
|  | H3K4me2 | 0.329 ± 0.015 | 0.313 ± 0.006 | **0.472 ± 0.008** | 0.338 ± 0.031 | 0.313 ± 0.005 | 0.347 ± 0.008 | <u>0.407 ± 0.006</u> |
|  | H3K4me3 | 0.361 ± 0.005 | 0.349 ± 0.029 | **0.466 ± 0.022** | 0.361 ± 0.024 | 0.403 ± 0.018 | 0.319 ± 0.006 | <u>0.424 ± 0.008</u> |
|  | H3K79me3 | 0.607 ± 0.013 | 0.602 ± 0.033 | <u>0.685 ± 0.023</u> | 0.611 ± 0.013 | 0.666 ± 0.008 | 0.612 ± 0.006 | **0.689 ± 0.013** |
|  | H3K9ac | 0.490 ± 0.029 | 0.545 ± 0.021 | **0.642 ± 0.010** | 0.550 ± 0.013 | 0.555 ± 0.005 | 0.541 ± 0.008 | <u>0.589 ± 0.005</u> |
|  | H4 | 0.742 ± 0.014 | 0.781 ± 0.004 | 0.789 ± 0.019 | 0.765 ± 0.024 | **0.851 ± 0.016** | 0.806 ± 0.005 | <u>0.823 ± 0.009</u> |
|  | H4ac | 0.376 ± 0.021 | 0.454 ± 0.024 | **0.574 ± 0.010** | 0.454 ± 0.008 | 0.496 ± 0.015 | 0.424 ± 0.004 | <u>0.500 ± 0.007</u> |
|  | Promoter All | 0.845 ± 0.020 | 0.825 ± 0.016 | 0.825 ± 0.025 | 0.847 ± 0.012 | **0.921 ± 0.011** | 0.897 ± 0.012 | <u>0.912 ± 0.005</u> |
|  | Promoter NoTATA | 0.886 ± 0.010 | 0.868 ± 0.025 | 0.920 ± 0.004 | 0.895 ± 0.008 | 0.913 ± 0.013 | <u>0.932 ± 0.017</u> | **0.960 ± 0.008** |
|  | Promoter TATA | 0.527 ± 0.032 | 0.650 ± 0.004 | 0.659 ± 0.006 | 0.600 ± 0.009 | 0.804 ± 0.016 | <u>0.813 ± 0.012</u> | **0.821 ± 0.003** |
|  | Splice All | 0.814 ± 0.018 | 0.858 ± 0.015 | 0.831 ± 0.012 | 0.815 ± 0.012 | <u>0.880 ± 0.025</u> | 0.836 ± 0.012 | **0.890 ± 0.003** |
|  | TF Human 1 | 0.630 ± 0.024 | 0.609 ± 0.020 | **0.719 ± 0.010** | 0.591 ± 0.017 | 0.644 ± 0.013 | 0.676 ± 0.009 | <u>0.680 ± 0.003</u> |
|  | TF Human 2 | 0.621 ± 0.013 | 0.679 ± 0.009 | 0.718 ± 0.013 | 0.640 ± 0.011 | 0.698 ± 0.013 | <u>0.724 ± 0.008</u> | **0.734 ± 0.013** |
|  | TF Human 3 | 0.597 ± 0.028 | 0.552 ± 0.013 | **0.634 ± 0.005** | 0.570 ± 0.016 | <u>0.621 ± 0.002</u> | 0.548 ± 0.006 | 0.611 ± 0.005 |
|  | TF Human 4 | 0.440 ± 0.011 | 0.521 ± 0.014 | **0.631 ± 0.006** | 0.474 ± 0.021 | 0.496 ± 0.023 | 0.499 ± 0.013 | <u>0.559 ± 0.011</u> |
|  | TF Human 5 | 0.700 ± 0.004 | 0.665 ± 0.002 | **0.789 ± 0.020** | 0.683 ± 0.024 | 0.710 ± 0.004 | 0.613 ± 0.002 | <u>0.760 ± 0.008</u> |
|  | TF Mouse 1 | 0.401 ± 0.005 | 0.540 ± 0.017 | 0.558 ± 0.009 | 0.494 ± 0.003 | **0.630 ± 0.011** | 0.466 ± 0.003 | <u>0.603 ± 0.009</u> |
|  | TF Mouse 2 | 0.840 ± 0.033 | 0.783 ± 0.005 | 0.848 ± 0.014 | 0.769 ± 0.016 | <u>0.859 ± 0.013</u> | 0.825 ± 0.015 | **0.870 ± 0.016** |
|  | TF Mouse 3 | 0.780 ± 0.021 | 0.684 ± 0.008 | <u>0.796 ± 0.017</u> | 0.686 ± 0.024 | 0.761 ± 0.015 | 0.707 ± 0.006 | **0.803 ± 0.009** |
|  | TF Mouse 4 | 0.585 ± 0.007 | <u>0.768 ± 0.018</u> | **0.771 ± 0.033** | 0.732 ± 0.014 | 0.667 ± 0.018 | 0.530 ± 0.005 | 0.704 ± 0.011 |
|  | TF Mouse 5 | 0.464 ± 0.004 | 0.413 ± 0.005 | <u>0.520 ± 0.013</u> | 0.423 ± 0.012 | 0.473 ± 0.026 | 0.423 ± 0.021 | **0.544 ± 0.010** |
| Nucleotide Transformer Tasks | Enhancer | 0.501 ± 0.022 | 0.448 ± 0.008 | 0.509 ± 0.011 | 0.500 ± 0.012 | **0.573 ± 0.008** | <u>0.537 ± 0.013</u> | 0.533 ± 0.004 |
|  | Enhancer Type | 0.498 ± 0.004 | 0.465 ± 0.005 | 0.528 ± 0.011 | 0.526 ± 0.013 | **0.576 ± 0.004** | <u>0.572 ± 0.017</u> | 0.553 ± 0.006 |
|  | H2AFZ | **0.535 ± 0.024** | 0.504 ± 0.031 | 0.482 ± 0.014 | **0.535 ± 0.022** | <u>0.534 ± 0.022</u> | 0.505 ± 0.008 | 0.516 ± 0.009 |
|  | H3K27ac | 0.458 ± 0.015 | 0.436 ± 0.011 | 0.497 ± 0.023 | <u>0.503 ± 0.015</u> | 0.434 ± 0.013 | 0.501 ± 0.008 | **0.531 ± 0.002** |
|  | H3K27me3 | 0.580 ± 0.025 | 0.553 ± 0.021 | 0.597 ± 0.018 | <u>0.612 ± 0.011</u> | 0.599 ± 0.006 | 0.588 ± 0.002 | **0.641 ± 0.010** |
|  | H3K36me3 | 0.578 ± 0.015 | 0.543 ± 0.014 | <u>0.651 ± 0.024</u> | 0.598 ± 0.006 | 0.592 ± 0.009 | 0.618 ± 0.009 | **0.667 ± 0.008** |
|  | H3K4me1 | 0.414 ± 0.004 | 0.386 ± 0.021 | 0.479 ± 0.011 | 0.389 ± 0.007 | <u>0.483 ± 0.004</u> | 0.481 ± 0.011 | **0.509 ± 0.003** |
|  | H3K4me2 | 0.508 ± 0.018 | 0.527 ± 0.024 | 0.501 ± 0.014 | 0.529 ± 0.004 | 0.549 ± 0.022 | <u>0.553 ± 0.008</u> | **0.585 ± 0.016** |
|  | H3K4me3 | <u>0.657 ± 0.007</u> | 0.606 ± 0.019 | **0.659 ± 0.005** | 0.638 ± 0.013 | 0.595 ± 0.022 | 0.629 ± 0.004 | 0.654 ± 0.003 |
|  | H3K9ac | 0.434 ± 0.006 | 0.490 ± 0.025 | 0.568 ± 0.019 | 0.513 ± 0.005 | <u>0.574 ± 0.011</u> | 0.538 ± 0.004 | **0.611 ± 0.015** |
|  | H3K9me3 | 0.406 ± 0.019 | 0.421 ± 0.011 | 0.463 ± 0.015 | 0.428 ± 0.023 | <u>0.485 ± 0.007</u> | <u>0.485 ± 0.013</u> | **0.520 ± 0.017** |
|  | H4K20me1 | 0.623 ± 0.016 | 0.590 ± 0.023 | 0.646 ± 0.017 | 0.656 ± 0.028 | 0.649 ± 0.001 | <u>0.659 ± 0.023</u> | **0.692 ± 0.004** |
|  | Promoter All | 0.693 ± 0.019 | 0.725 ± 0.029 | 0.724 ± 0.013 | 0.709 ± 0.007 | **0.788 ± 0.008** | 0.760 ± 0.012 | <u>0.787 ± 0.004</u> |
|  | Promoter NoTATA | 0.726 ± 0.013 | 0.737 ± 0.013 | 0.738 ± 0.020 | 0.762 ± 0.012 | **0.823 ± 0.005** | 0.767 ± 0.013 | <u>0.807 ± 0.009</u> |
|  | Promoter TATA | 0.840 ± 0.004 | 0.798 ± 0.011 | 0.832 ± 0.010 | 0.830 ± 0.007 | 0.866 ± 0.009 | <u>0.905 ± 0.005</u> | **0.960 ± 0.009** |
|  | Splice Acceptor | 0.783 ± 0.003 | 0.844 ± 0.021 | 0.794 ± 0.022 | 0.836 ± 0.022 | <u>0.962 ± 0.004</u> | 0.959 ± 0.007 | **0.971 ± 0.006** |
|  | Splice All | 0.843 ± 0.014 | 0.926 ± 0.013 | 0.828 ± 0.031 | 0.847 ± 0.011 | **0.971 ± 0.014** | 0.953 ± 0.007 | <u>0.969 ± 0.006</u> |
|  | Splice Donor | 0.884 ± 0.014 | 0.952 ± 0.019 | 0.808 ± 0.006 | 0.831 ± 0.003 | <u>0.972 ± 0.019</u> | <u>0.972 ± 0.002</u> | **0.984 ± 0.011** |
| Avg. Score |  | 0.612 | 0.632 | 0.674 | 0.626 | <u>0.684</u> | 0.654 | **0.704** |
| Avg. Rank |  | 5.836 | 5.309 | 3.327 | 5.009 | <u>2.800</u> | 3.936 | **1.782** |

